# DNA origami uptake in Y-79 retinoblastoma cells driven by oligolysine coating

**DOI:** 10.64898/2026.06.08.730913

**Authors:** Anna Klose, Zahra Gounani, Sergei Raik, Artturi Koivuniemi, Sonja Korhonen, Mika Reinisalo, Tatu Lajunen, Veikko Linko, Timo Laaksonen

## Abstract

DNA origami nanoparticles (DONs) are attractive nanocarriers of controllable size, shape and addressability that have potential for treating eye diseases by overcoming ocular barriers. However, suboptimal physiological stability and poor cell uptake due to the negative charge may limit their use. Previous reports show that electrostatic complexation of DONs with cationic PEG-oligolysine block-copolymers like PEG5K-K10 can improve structural integrity and promote cell internalization. Here, we investigated a dual approach of PEG5K-K10 coatings and PL3 targeting peptides to improve uptake of 24-helix bundle (24HB) DONs into Y-79 retinoblastoma cells. Uptake studies revealed that PEG5K-K10 was essential for DON uptake in Y-79 cells, as uptake only occurred upon exceeding a distinct PEG5K-K10 amount. Longer exposure times or increased polymer amounts improved cell association. However, no beneficial effect of PL3 was observed. While free PEG5K-K10 reduced cell viability at higher concentrations (IC_50_ 36.8 µM), coated DONs were well-tolerated. Furthermore, single particle tracking in ex vivo porcine eyes revealed comparable vitreal mobility for uncoated and coated 24HB, with a slight decrease at higher coating amounts. Our findings highlight that PEG5K-K10 can enhance ocular cell uptake without limiting nanoparticle diffusivity in the vitreous, and support further optimization of DONs for ocular drug delivery.

## Introduction

DNA origami nanoparticles (DONs) are promising nanocarriers in therapeutic applications, offering a versatile and precise platform for modulating their size, shape and functionalization.^1–3^ Through sequence-specific DNA hybridization between a long single-stranded DNA (ssDNA) scaffold strand and short, base-complementary oligonucleotides (staple strands), DONs of defined 2D and 3D structures are folded and can be decorated in a spatially-controlled manner with various targeting moieties.^4–7^ Often limited in their stability in vitro and in vivo due to low salt strength and nuclease susceptibility,^8–10^ modulating staple strand cross-over spacing,^11^ ligating DNA backbone breaks,^12–15^ UV-crosslinking of the DON structure,^16,17^ or electrostatically assembled, protective coatings of proteins,^18–20^ lipids,^21–23^ and polymers^24–29^ offer suitable mitigation strategies.

Especially commercially available block-copolymers of oligolysine (K10) with polyethylene glycol units (PEG1K or 5K), referred to in the following as PEG1K-K10 and PEG5K-K10, have been extensively studied, contributing to structural and colloidal stability, and improved cell uptake of DONs.^24,25,30–34^ While DONs by themselves may often display only limited cell internalization, the polymeric coating can mask their negative charge, and the coating amount can be adjusted to the preferences of respective cell lines.^34^ Furthermore, PEG-K10 coatings leave functionalities of the underlying DON addressable^25,31^ and have been successfully utilized in combination with targeting ligands such as peptides,^25^ aptamers,^30^ antibodies.^29^ The coating can also reduce unspecific cell binding,^29,35^ though some concerns regarding the binding capacity of targeting moieties to specific receptors due to steric effects of the PEG-chains have been reported.^36,37^

Initially, DONs were investigated mainly for cancer-related applications.^38^ Recently though, interest is also growing in the application of DONs for e.g., vaccine and immune therapies *via* specific templating of stimulatory ligands,^39–41^ and for inflammatory pathologies like rheumatic arthritis, kidney and lung conditions due to their intrinsic ROS scavenging capabilities.^42–44^ Similarly, a recent report from Wu *et al.* demonstrated that the targeted combination therapy of anti-VEGF (vascular endothelial growth factor) aptamers and antibodies on a DNA origami rectangle synergistically improved choroidal neovascularization (CNV) lesions in murine eyes by local reduction of VEGF activity and oxidative stress.^45^ These results offer a promising outlook for the application of DONs in ocular diseases like age-related macular degeneration (AMD) and diabetic retinopathy, that despite different pathogenesis, result in progressive inflammatory conditions. They are characterized by elevated reactive oxygen species’ levels and aberrant vessel formation, ultimately leading to vision loss.^46–49^

Impaired vision and blindness impose substantial burden on patients and healthcare systems and require improved treatment strategies.^50–53^ One of the main challenges in treating conditions affecting the posterior segment of the eye is ensuring intraocular bioavailability and therapeutic concentrations which are limited in systemic administration by the ocular barrier tissues.^54^ Nanoparticles offer an attractive platform to tailor their properties for overcoming such barriers and increasing intraocular retention.^55,56^ DONs, compatible with a variety of therapeutic molecules, could be harnessed to deliver combination therapies in a targeted manner. To deliver therapeutics to the retina, the common administration route is local intravitreal injection into the eye, bypassing the blood-ocular barrier and achieving higher intraocular concentrations. The vitreous humor is a gel-like matrix with a mesh size of ∼500 nm,^57^ that fills out the inner eye and consists of water (>98%), collagen and negatively charged glycosaminoglycans that serve as a size-and charge-dependent diffusion barrier for drugs and nanoparticles.^58,59^ Due to their inherent negative charge and defined size, we would expect DONs to remain mobile in the vitreous, and thus, verifying their mobility is an important step in the development of intravitreal DONs. In our previous work, we observed that DONs achieved high drug loadings of the chemotherapeutic drug doxorubicin and curbed its cell toxicity, but the low nanoparticle uptake into ARPE-19 retinal cells limited its effectiveness as drug nanocarrier.^60^ Retinoblastoma is a rare but the most common pediatric intraocular tumor.^61^ Despite the high survival rate in developed countries, more efficient treatment options are needed for preserving vision and reducing the adverse effects on affected children. The therapeutic efficacy could be improved with targeted, nanoparticle treatments.^62^ Hence in this work, we aimed to investigate two strategies to improve selective cell internalization of 24-helix bundle (24HB) DONs into Y-79 retinoblastoma cells by (i) coating 24HB with PEG5K-K10 polymer and by (ii) incorporation of the targeting peptide PL3.

The PL3 peptide (AGRGRLVR) belongs to a subclass of tumor homing peptides,^63^ that display an exposed sequence motif of [R/K]XX[R/K] (X representing any amino acid) at their C-termini (“C-end Rule” (CendR) motif).^64^ Their free carboxy group can then bind to the cell surface receptor neuropilin-1 (NRP-1), which plays an important role in the development of nervous and vascular systems, thus also widely expressed in blood vessels of e.g. tumors, or in eyes during CNV.^65–68^ PL3 also targets the C-isoform of tenascin-C (TNC-C), an extracellular matrix glycoprotein, that is similarly upregulated in cancerous tissue and under inflammatory conditions like CNV.^63,69–71^ CendR targeting peptides on nanoparticles enhance tissue penetration and drive cellular internalization through NRP-1-mediated endocytosis.^64,72,73^ For instance, PL3-coated, PEGylated silver nanoparticles and iron oxide worms showed improved cell uptake and targeting of carcinoma tissue in mice.^63^ Overall, PL3 is a promising targeting moiety for ocular diseases since intravitreally injected PL3 peptide by itself accumulated in murine CNV sites, reducing angiogenesis and vascular leakage.^74^ Furthermore, PL3 demonstrated specific internalization and nucleolar accumulation in Y-79 retinoblastoma cells, expressing NRP-1 and TNC-C receptors.^75^

Here, we demonstrated that PL3-oligonucleotides could be attached via DNA hybridization to 24HB DONs and subsequently coated with PEG5K-K10 polymer. We studied the uptake of coated 24HB in Y-79 retinoblastoma cells depending on the PEG5K-K10 amount and the targeting peptide. Our findings revealed a clear uptake preference of Y-79 cells for coated DONs upon exceeding a certain PEG5K-K10 amount but could not resolve the benefit of the targeting peptides for uptake or cell association. Further investigations will be necessary to identify and improve the targeting properties of our DONs in that regard. Overall, the PEG5K-K10 coating enabled the uptake of DONs in ocular cells in vitro, with little toxicity. Lastly, we investigated the vitreal mobility of uncoated and coated 24HB DONs in ex vivo porcine eyes with single-particle tracking, mimicking future intravitreal injections. While the fraction of immobilized nanoparticles seemed to increase with higher coating ratios, we observed mostly similar diffusivity between the different 24HB samples and concluded that the necessity of cationic coatings for DONs could still be compatible with future intravitreal applications.

## Results and Discussion

### Characterization of fluorophore- and peptide-labeled 24HB DNA origami nanoparticles

In retinal ARPE-19 cells, only limited uptake was previously observable for 24-helix bundle (24HB) DNA origami nanoparticles (DONs), possibly due to lacking recognition by specific endocytic uptake pathways or due to their unfavorable, inherent highly negative charge.^60^ Hence, in this work, we probed whether the uptake of 24HB DONs into an ocular cell line could be modulated and improved by incorporating the targeting peptide PL3, and by modifying the apparent surface charge of 24HB with a cationic polymeric coating.

PL3 peptides (sequence: AGRGRLVR) effectively internalize and accumulate in Y-79 retinoblastoma cells,^75^ and could, thus, also facilitate the uptake of 24HB. To attach the targeting peptides, the 24HB structure^76^ was modified by extending single-stranded (ss)DNA overhang strands from the end of the DON that could hybridize with complementary DNA strands linked to the targeting peptide PL3 (Supporting information (SI) Section SI1). Depending on the experiment, four different versions of 24HB were utilized (Figure 1a): A regular 24HB DON without any modifications (not shown here),^76^ a fluorescently labeled 24HB (24HB-F),^60^ and a fluorescently labeled and PL3-conjugated 24HB (24HB-RFLP). 24HB-RFLP was also prepared when needed as a control without peptide attachment (24HB-RF). Similar to the peptides, Atto488 fluorophores were incorporated *via* extending DNA overhang strands of specific sequence (Table SI1),^60^ allowing for 24 attachments on 24HB-F. In case of 24HB-RFLP, the attachment sites for PL3 and Atto488 numbered each 12 and were separated to different ends of the structure to avoid any interference. While the incorporation of the Atto488 fluorophore and the PL3 peptide into 24HB-RFLP was not visible under the transmission electron microscope (TEM, Figure 1b), it led to observable changes during agarose gel electrophoresis (AGE, Figure 1c). The addition of the PL3 peptide resulted in a slight band shift for 24HB-RFLP compared to 24HB-RF due to size increase. The fluorescent signal in the Atto488 imaging channel corresponded with the bands of the fluorescent DONs.

**Figure 1.**
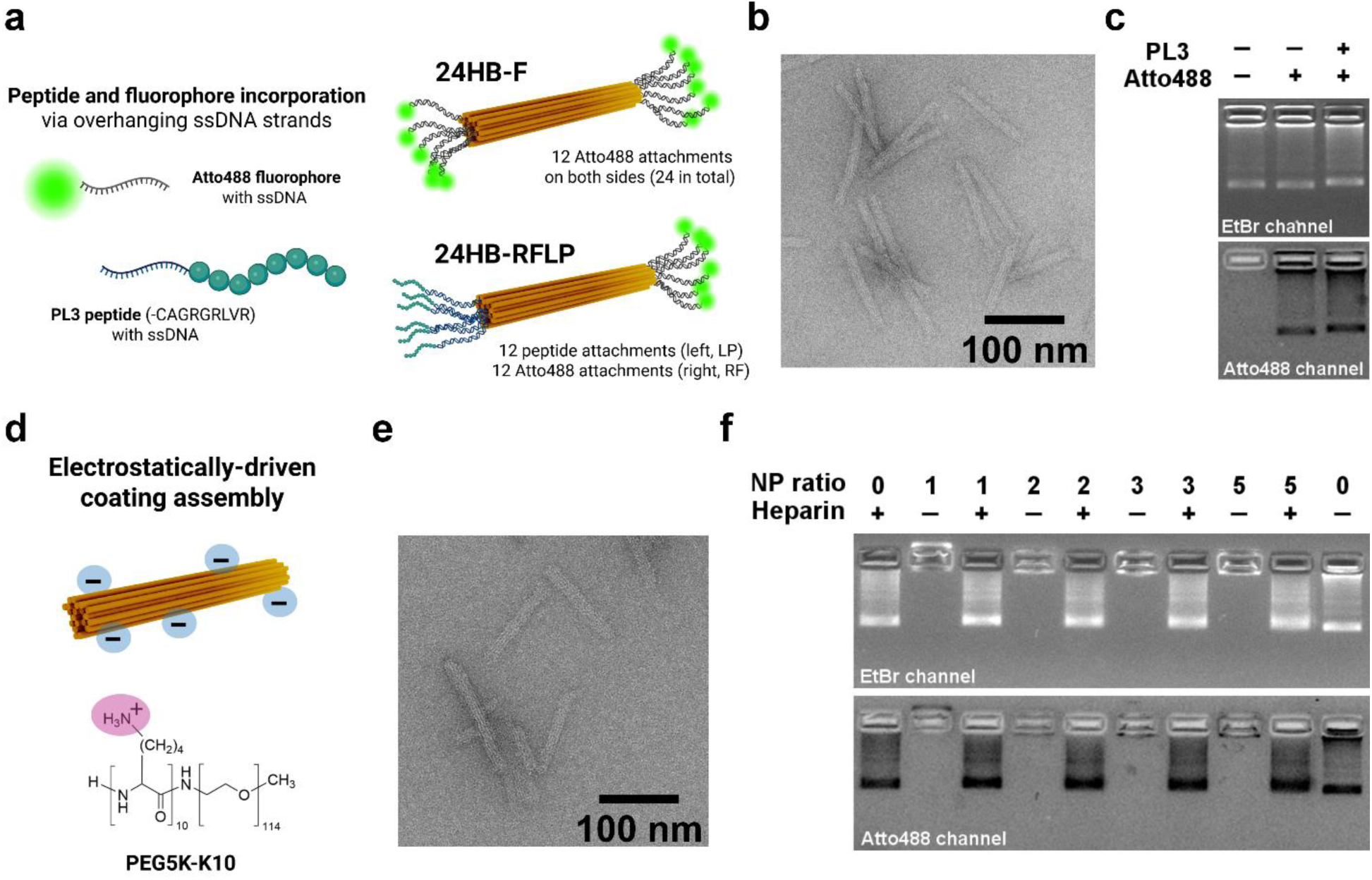
Overview and characterization of functionalized 24HB DONs in this work. **a.** Schematic representation of Atto488 fluorophore- and PL3 peptide-modified 24HB. Atto488- and PL3-functionalized oligonucleotides were annealed specifically to sequence-complementary single-stranded (ss)DNA overhangs from the 24HB. In this work, unmodified, regular 24HB (not shown here), fluorescent 24HB-F (top), and fluorescent and PL3-labeled 24HB-RFLP (bottom) were utilized. Note, 24HB-RFLP was prepared when needed also as its peptide-free control (24HB-RF, ssDNA overhangs for PL3 left unhybridized). **b**. TEM image of 24HB-RFLP. **c.** Incorporation of Atto488- and PL3-strands into the 24HB structure, visualized *via* agarose gel electrophoresis (AGE). 24HB with unhybridized overhangs for PL3 and Atto488 (-/-), 24HB-RF with unhybridized overhangs for PL3 (-/+) and 24HB-RFLP (+/+) appeared as visible bands in the ethidium bromide (EtBr) channel, and if fluorescent, in the Atto488 channel. The band shift of 24HB-RFLP indicated size increase due to PL3 attachment. **d.** Electrostatic coating assembly between positively charged lysines residues (N) in PEG5K-K10 polymer and negatively charged phosphate (P) groups in 24HB at distinct N-to-P charge ratios (NP). **e.** TEM image of coated 24HB-RFLP at NP2 ratio. **f.** AGE of uncoated and coated 24HB-F at different NP coating ratios (NP0-5). PEG5K-K10-coated 24HB-F became immobilized in the gel pocket, but upon addition of negatively charged heparin in 50-fold negative charge excess to PEG5K-K10, coatings were removed and similar bands to the uncoated reference (NP0) were visible in the EtBr and Atto488 channel. AGE images were cropped as needed. TEM samples in b and e were negatively stained with 2% uranyl formate and selected images cropped.

To investigate the effect of a cationic coating on cellular uptake of 24HB, PEG5K-K10 was electrostatically assembled onto 24HB DONs in 1× folding buffer (1×FOB) by simple mixing of the two components (Figure 1d).^25^ By varying the relative quantities of polymer to DONs, different charge ratios, expressed as N-to-P ratio (NP), were achievable. These NP ratios reflect the number of positive charges from the protonated amine groups (N) in the lysines and the negative charges provided by the phosphate groups (P) in the DONs, meaning a coating ratio of NP0 corresponded to an uncoated 24HB, while NP2 was prepared at a two-fold positive charge excess (Section SI2). At coating ratios between NP1 and NP5, PEG5K-K10 coatings on 24HB lead to no large visible changes or structural deformations under TEM (Figure 1e, Figure SI1). This is in line with previous reports, that the shape of coated DONs is dictated by the underlying DON structure and the PEG units support colloidal stability.^25,29,31,33,77^ On AGE, coated DONs got instantly immobilized in the gel pocket, as previously reported (Figure 1f).^25^ By incubating the coated DONs with heparin (Figure SI2), their coating was removed and their electrophoretic mobility regained, as verified in the ethidium bromide (EtBr) and Atto488 channel. While PEG5K-K10 coatings stabilized 24HB DONs against degradation and fluorophore detachment in cell media (Figure SI3),^24,25^ subsequent cell treatments with DONs were carried out in FBS-free media to limit the fluorophore loss for uncoated DONs and reduce general fluorescence background noise.

### Uptake studies in Y-79 cells

To study the effect of the PL3 targeting peptide and the PEG5K-K10 coating on DON uptake, Y-79 cells were exposed for 24 h to uncoated and NP2-coated 24HB-RF and 24HB-RFLP. Based on the overlay of the fluorescence intensity histograms, it became apparent that there was only a minor shift for uncoated 24HB-RF (pink) and 24HB-RFLP (red) towards higher fluorescence intensities compared to the buffer-treated control (beige), indicating that Y-79 cells did not internalize much uncoated DONs (Figure 2a). In contrast, the change in shape and shift of the fluorescence intensity histograms for coated NP2-24HB-RF (green) and NP2-24HB-RFLP (blue) suggested that the coating promoted DON uptake.

**Figure 2.**
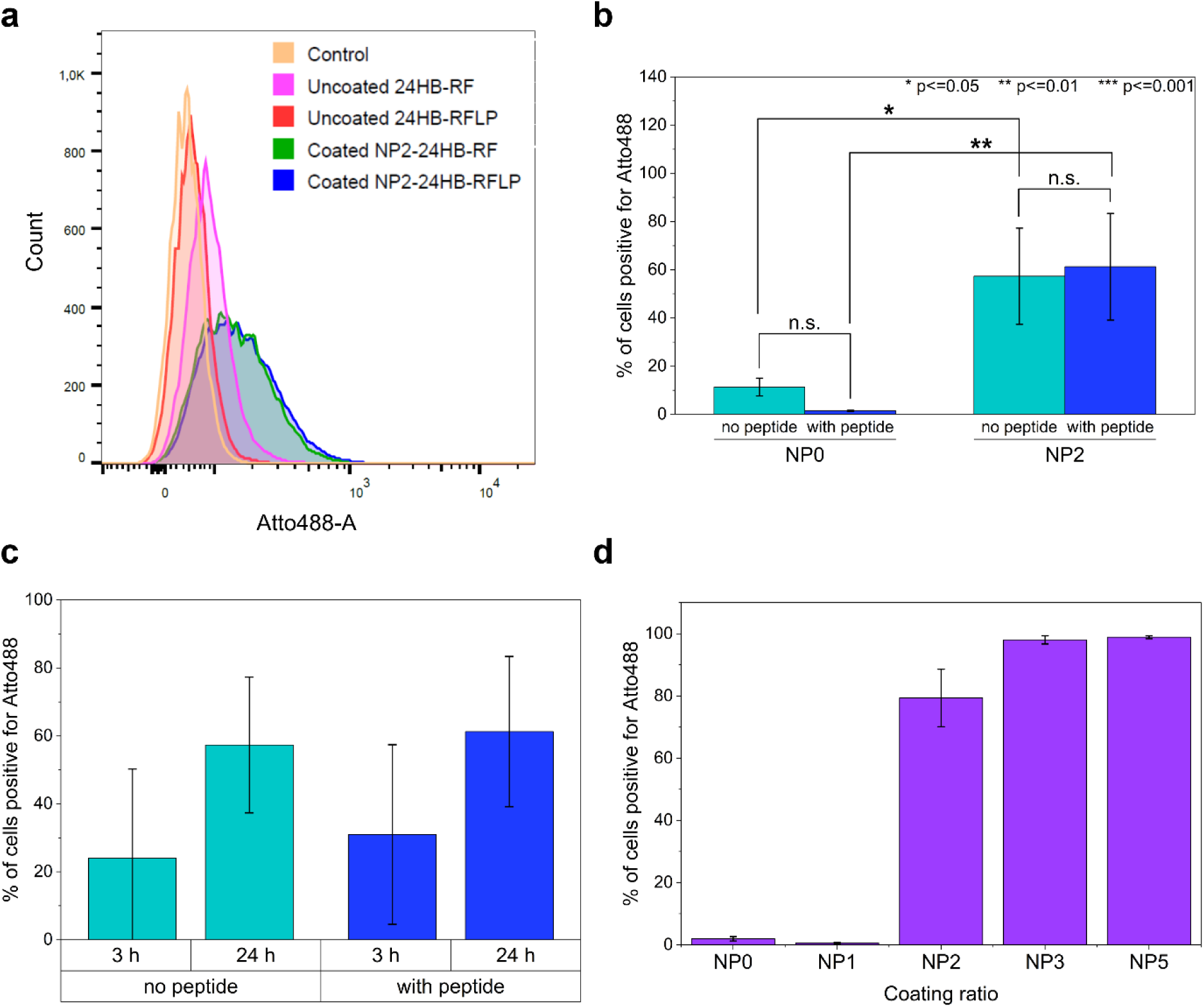
Uptake of uncoated and coated 24HB-RF (no peptide) and 24HB-RFLP (with peptide) in Y-79 cells with flow cytometry. Uncoated (NP0) and coated (NP2) DONs were incubated at 1.5 nM final concentration with cells in RPMI-1640 media with 1% PS. **a.** Exemplary histogram of cell fluorescence in the Atto488 channel after 24 h. **b.** Percentages of treated cells positive for Atto488 after 24 h, depending on the presence of the coating (NP0 vs. NP2) and the targeting peptide. **c**. Percentages of cells positive for Atto488 that were treated with coated DONs at coating ratio NP2 for either 3 h or 24 h. **d**. Percentages of cells positive for Atto488 that were treated for 24 h with coated 24HB-RFLP at different coating ratios (NP0-5). Mean ± s.d. was reported, acquired from three independent experiments (n=3) with two technical replicates each (for b and c). In case of d, the data was collected in several independent experiments (n=2-4, n=2 for NP1 and NP5), with one to two technical replicates each. * p ≤ 0.05, ** p ≤ 0.01, *** p ≤ 0.001, n.s. (not significant). 2-way ANOVA followed by Tukey post-hoc test was used for data analysis.

This observation was also supported by flow cytometry analysis when considering the percentages of cells positive for the Atto488 signal (Figure 2b, Table SI2): Uncoated (NP0) 24HB-RF and 24HB-RFLP were taken up on average by only 11.4% and 1.5% of the cells respectively, underlining that the PL3 peptide alone could not promote the DON uptake. Possibly their inherent negative charge hindered their internalization, yet it was rather unexpected that the peptide-free control led to more cell association, since the peptide was supposed to promote receptor binding. On the other hand, coated 24HB-RF and 24HB-RFLP at the coating ratio of NP2 led to 57.3% and 61.3% of cells positive for Atto488, respectively, demonstrating that the coating enhanced the DON uptake in Y-79 cells, as observed previously in other cell lines.^25,34^ This was also underscored by the result of the two-way ANOVA, revealing that the percentage of cells positive for Atto488 was significantly affected by the presence of the coating (*F*(1,8) = 37.192, p = 0.0003). Post-hoc comparisons using Tukey HSD test confirmed that the increase in the percentage of cells positive for Atto488 for both NP2-24HB-RF and NP2-24HB-RFLP was significant compared to their corresponding uncoated counterpart (p = 0.0234, p = 0.0054). The incorporation of the peptide, however, showed no significant effect (*F*(1,8) = 0.118, p = 0.740), neither was the interaction between the peptide and the coating significant (*F*(1,8) = 0.635, p = 0.449). Overall, there was no additional benefit of the incorporated peptide observed under these conditions, however, the coating enhanced the DON uptake, which was also verified in ARPE-19 cells (Figure SI4).

Additionally, the uptake of coated NP2-24HB-RF and NP2-24HB-RFLP was compared after 3 h and 24 h, revealing that the longer incubation promoted DON uptake by more Y-79 cells, regardless of the peptide (Figure 2c). During replication of the measurements, the absolute percentages of cells positive for Atto488 varied across different days, resulting in the higher standard deviation shown. Yet, within a set of measurements from the same day, we observed consistently that the 24 h incubation outperformed a 3 h incubation (Table SI2). Similar increase in cell uptake and cell attachment with prolonged incubation time has been observed previously, though specific uptake kinetics and extent of accumulation seemed cell line dependent.^33,78^ Remarkably, increasing the coating ratio above NP2 resulted in an Atto488-signal by almost all cells (NP3: 98.0%, NP5: 98.9%; Figure 2d). For coating ratios NP0 and NP1, uptake was very limited and negligible (NP0: 2.0%, NP1: 0.6%). A narrowly spaced-out screening for NP ratios between 0 and 2.5 suggested that exceeding a distinct threshold of minimum NP1.75 was necessary to promote uptake in Y-79 cells (Figure SI5). This was unexpected as most PEG-oligolysine coated DONs are usually studied at coating ratios NP1,^25,33^ but it is in line with the systematic study of Koga *et al.* that different cells exhibit different uptake preferences regarding the surface charge of DONs.^34^ To summarize, uncoated 24HB DONs were not well internalized in Y-79 cells, similar to ARPE-19 cells.^60^ The incorporation of PL3 peptide into 24HB-RFLP resulted in no enhanced uptake over 24HB-RF, neither in an uncoated or coated state. Coating of 24HB DONs with PEG5K-K10 polymer at a coating ratio of NP2, however, increased cell uptake over time, with even higher coating ratios leading to more cell association.

While targeting ligands on DNA nanostructures do not necessarily always enhance their performance,^79^ cationic targeting peptides like cell penetrating peptides ^42,80,81^ or cyclo[Arg-Gly-Asp-(D-Phe)-Cys] (cRGD) peptides that interact with specific αvβ3 integrins have enhanced cell internalization of DONs previously.^82,83^ Cend peptides are particularly promising for similar application, as they have demonstrated micropinocytosis-like, receptor-mediated internalization through formation of large endocytic vesicles that could even accommodate payloads that were not covalently attached.^72,84^ In line, PEGylated and PL3-decorated metal nanoparticles demonstrated cell uptake and cancer targeting in vivo.^63^ PL3 peptides were specifically selected as our ocular targeting ligands since the peptides by themselves demonstrated rapid cell binding and internalization in Y-79 retinoblastoma cells.^75^ While we believe that the positioning of the targeting ligand at the end of the DON rod should have facilitated axial receptor binding,^36^ it is possible that our locally concentrated arrangement of all PL3-peptides on one end was less favorable for the ligand-receptor interaction. By modulating ligand density and spatial distribution,^85^ e.g. by spacing a higher number of ligands more evenly along the entire 24HB structure, we could perhaps improve receptor engagement and promote a favorable orientation for membrane binding and engulfment.^37,86^ More compact DONs shapes are generally preferred for cell uptake and should be prioritized to maximize effectiveness in the future,^33^ though we demonstrated that even DONs of elongated shape like 24HB can enter cells.^86^ Refining our experimental conditions, inclusion of fetal bovine serum, and addition of proteins that bias the protein corona favorably could further improve cell uptake.^32^ If combined with additional insights into the distribution, expression behavior and pattern of the target receptor in Y-79 cells, DONs could still hold great potential for optimized spatial presentation of the PL3 peptide, as already realized for other ligands in immunomodulatory applications.^39,41^

Generally, electrostatic DNA-peptide interaction is the basis for the assembly of our PEG5K-oligolysine coating onto DONs.^25,29,87^ This also raises the question though, whether the positive charge of our targeting peptide could negatively affect its incorporation and display on DONs. For instance, Kang *et al.* reported that cationic peptides can unselectively adsorb onto the DON surface, resulting in uncontrolled loading stoichiometry and necessitating several repeated purification cycles. Most importantly though, they showed for PEG5K-K10 coated, peptide-decorated DON at NP1, that only site-specifically hybridized peptides elicit a biological response through receptor interaction.^78^ While some of the cationic peptides examined in their work have a similar net charge at pH 7 to PL3 (+3), none of them displays a Cend motif, and moreover, PL3 differs in length, sequence, molecular weight and has an increased isoelectric point (pI 12.2). Our molecular dynamics simulation of a free dsDNA molecule and PL3 peptides showed that PL3 strongly interacts and forms stable complexes with DNA (Figure SI6). However, since the incubation of PL3-oligonucleotides with 24HB DONs without specific ssDNA overhangs led to no observable band shift on AGE (Figure SI7), the extent of unspecific PL3 adsorption appeared to be limited in our case or at least easily stripped off in the electrophoretic field applied during AGE, suggesting incorporation of PL3 *via* hybridized linker strands in 24HB-RFLP. Furthermore, our dsDNA spacer of 23 nucleotides between PL3 and 24HB is of comparable length to other works,^31,42,78,82^ and considering the persistence length of dsDNA (∼150 bp), the spacer should offer some rigidity in displaying the targeting peptide. Possibly though, the conformational display and receptor-binding of PL3 is affected by the proximity of the DNA linker, the DONs structure itself, or obscured by our PEG5K-K10 coating due to steric effects.

There are some reports indicating that the PEG-oligolysine coatings can reduce receptor binding.^36,37^ On the other hand though, many reports also demonstrated accessibility of ssDNA strands after coating with PEG1K-K10 and PEG5K-K10 and their compatibility with a range of different attachments, including targeting ligands.^24,25,88^ For instance, functionalization of a DNA origami rod with cRGD peptides and subsequent coating with PEG5K-K10 demonstrated successful targeting against the surface integrin receptor and enhanced cell uptake compared to an untargeted, coated DON.^25^ Similarly, aptamer-decorated, PEG5K-K10-coated DONs at NP 1 outperformed the non-targeted control in a mouse inflammation model.^30^

Ponnuswamy *et al.* reported that the attachment of PEG5K at NP1 increased DON thickness by 2.4±1.3 nm according to dynamic light scattering.^25^ The homogeneity of PEG-oligolysine coatings and their effect on attachments is not fully resolved yet, though coatings seem to be at least present in both central and edge locations of DONs.^89^ Given the 23 bp length (∼7.8 nm) of our PL3-oligonucleotide linker, we expected our peptide attachment to extend out of the coating and be accessible, especially due to their position on the outer edge of the DON, which according to digestion studies is a more accessible position under the protective coating.^31^ At the same time though, 21 bp long DNA extensions were sufficiently protected against nucleases by even only PEG1K-K10 coatings at the NP ratio of 0.75,^31^ underlining that even PEG1K layers exert some steric effects which could then be even more pronounced for PEG5K-K10. Furthermore, higher coating ratios like we used in this work would result in denser coverage of the DON surface, leading to a denser PEG brush conformation that would likely increase also the PEG layer thickness (Section SI11). In most investigations with PEG-oligolysine coated DONs thus far, however, they have only focused on one specific NP ratio (usually ≤ NP1). In contrast, Cremers *et al.* reported that PEG5K-K10 coating at the NP ratio of 2.5 decreased the cell surface accessibility and receptor binding compared to an uncoated, antibody-targeted DON rod.^36^ Thus, further clarification is warranted on how exactly ligand accessibility scales with increasing NP ratios and different targeting ligands. At this point, our results suggest that PEG5K-K10 coating is the main driving force for uptake into Y-79 cells, most likely through non-specific electrostatic interactions. Overall, this underlines the need for further investigations into the receptor binding capacity of coated, peptide-functionalized 24HB in cell-free and cell environments to discern and resolve the contribution and effect of the coating and the targeting ligand. That would in turn allow further optimization of experimental, biological and DON related conditions and factors, that could improve targeted cell uptake.

Our findings from the flow cytometry studies were also supported by live confocal imaging: Adherently grown Y-79 cells were treated for 24 h with uncoated 24HB-RFLP or NP2-coated 24HB-RFLP (Figure 3). Free PL3 peptide has previously demonstrated nucleolar accumulation,^75^ but this was not observable for our DONs here. While uncoated 24HB-RFLP seemed to only enter and bind dead cells (green signal in Atto488 channel), the signal for NP2-24HB-RFLP colocalized with the signal for LysoTracker Deep Red (red; combined: yellow), indicating that coated DONs were present in lysosomes.^25^ The uptake of NP2-24HB-RF showed a similar pattern (Figure SI8). Likewise, coated DONs were also internalized into ARPE-19 cells, though they accumulated only a little along the outer cell membranes, and showed some colocalization with Lysotracker and possibly distribution into the cytosol (Figure SI9). In Y-79 cells, NP2-24HB-RFLP seemed to display rather some affinity for the outer cell membranes, as the fluorescent signal from the DONs clearly outlined the cells.

**Figure 3.**
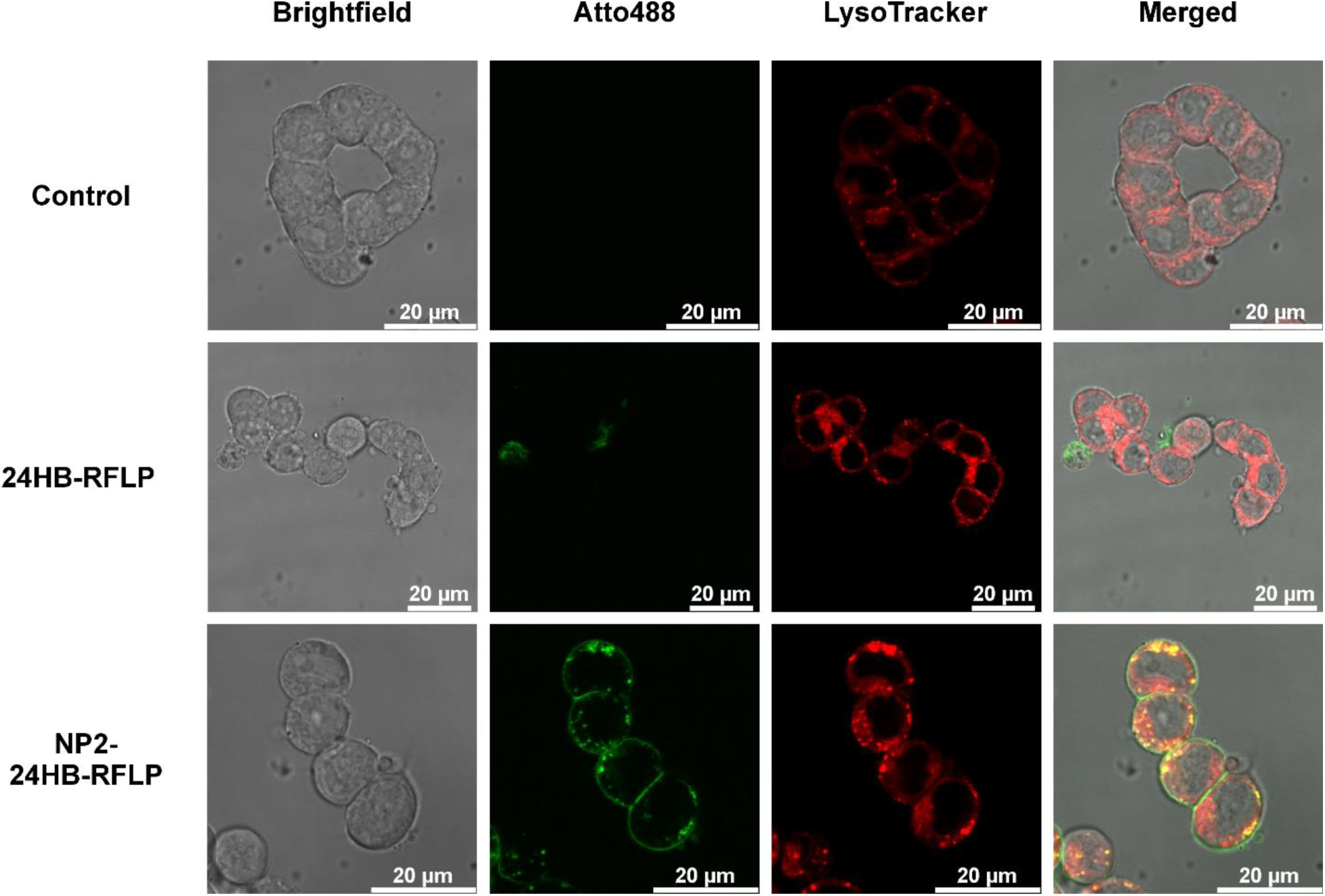
Live confocal imaging of Y-79 cells. Cells were treated with uncoated and coated NP2-24HB-RFLP for 24 h at 10 nM final concentration in RPMI-1640 media with 1% PS. 24HB-RFLP was functionalized with Atto488 fluorophores (green), lysosomes in cells were stained with LysoTracker Deep Red (red), buffer-treated cells served as control. Colocalization of Atto488 and LysoTracker Deep Red resulted in the yellow signal in the merged images.

Possibly, electrostatic interactions between the coated DONs and the Y-79 cell surface promoted cell association, which could be therapeutically beneficial. This was not resolvable in flow cytometry, where associated and internalized DONs alike contribute to the fluorescent signal.^33^ Another explanation is the ∼6.7 fold higher DONs concentration used for the visualization in confocal imaging that could contribute to some overcrowding and might not necessarily mirror our flow cytometry uptake studies at lower DON concentration.^34^ Unspecific cell binding of PEG5K-K10 block-copolymer might be, however, to a certain extent unavoidable. Extensive investigation of PEG-grafted oligolysines elucidated relevant design parameters to further optimize DON coating and performance, surpassing the commercial PEG5K-K10 block-copolymer that was used in this work.^29^ For instance, PEG-oligolysine coatings could reduce specific and unspecific cell binding in dendritic cells, especially with increasing NP ratios but to varying extent depending on the polymer.^29^ In contrast, antibody-decorated, commercial PEG5K-K10-coated DONs at NP 0.5 showed little cell binding specificity compared to the non-targeted control.^29^ The commercial availability has facilitated however its wide-spread use as a coating agent for DONs, and while perhaps not optimal, has effectively promoted the interaction and uptake of our PEG5K-K10 coated 24HB DONs into ocular cells.

### Cell viability assessment of PEG5K-K10 polymer and coated DONs

As beneficial as cationic polymers are for enhancing nanoparticle uptake, depending on the concentration of free polymer, they could display membrane-disruptive and toxic properties against cells.^90^ To assess the properties of free PEG5K-K10 polymer, Y-79 cells in suspension were treated in FBS-containing cell media for 24 h, before their metabolic activity was analyzed *via* alamarBlue assay and compared to the buffer-treated control, that was set to 100% viability (Figure 4a). With increasing concentration of free PEG5K-K10, the cell viability of Y-79 cells decreased. This effect would appear to be, however, also cell-line specific, as an exploratory screening of free PEG5K-K10 in the same concentration range impacted ARPE-19 cells far less (Figure SI10). For Y-79 cells, the IC_50_ value for free PEG5K-K10 under these conditions was 36.8 ±7.6 µM.

**Figure 4.**
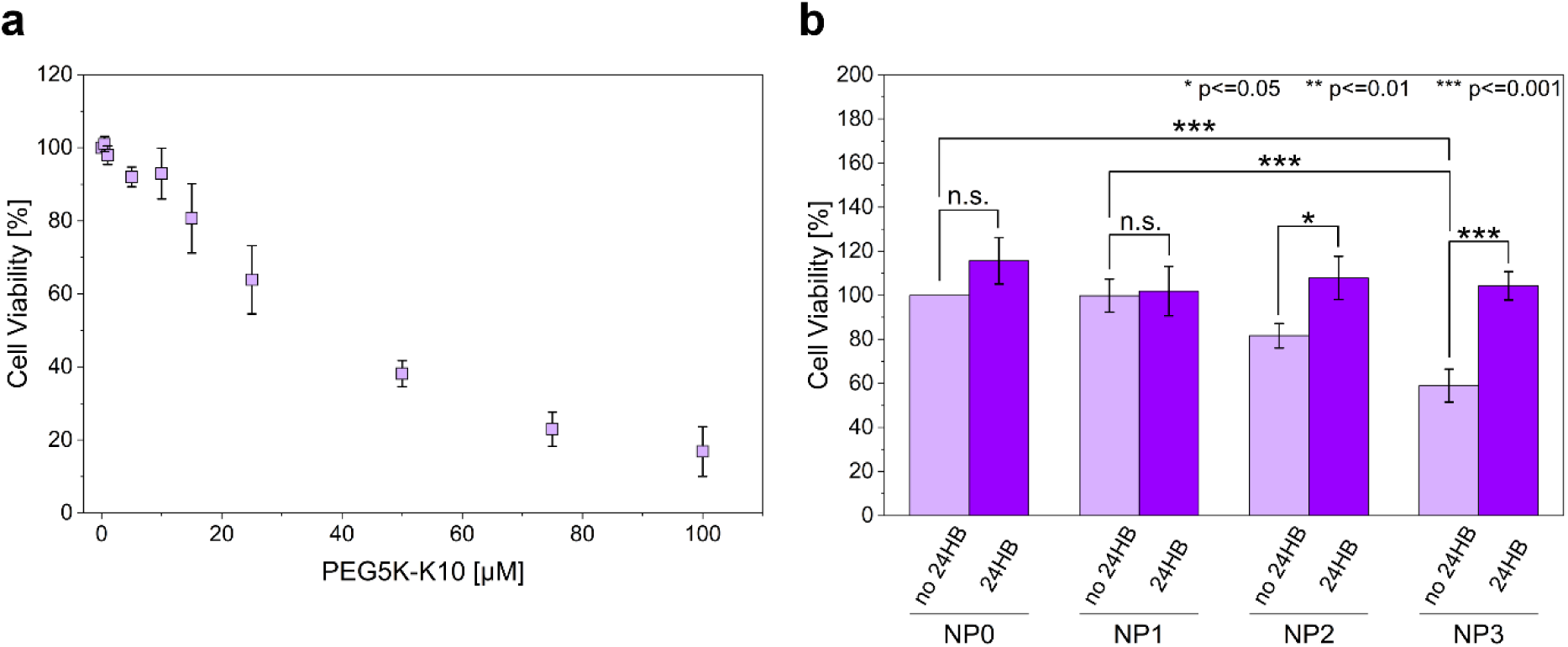
Cell viability of Y-79 cells after 24 h exposure, assessed *via* alamarBlue assay. Cells were treated with **a.** free PEG5K-K10 polymer (0.5-100 µM), **b.** uncoated (NP0) and coated 24HB (2 nM) at different coating ratios (NP1-3) and controls of corresponding amounts of free polymer (no 24HB). Y-79 cells in **a** were grown in suspension and exposed under FBS-containing conditions, while cells in **b** were grown adherently and treated under FBS-free conditions, in correspondence to the uptake experiments. Buffer-treated cells served as control (set to 100% viability). Mean ± s.d. was reported, acquired from three independent experiments (n=3) with each three technical replicates. * p ≤ 0.05, ** p ≤ 0.01, *** p ≤ 0.001, n.s. (not significant). 2-way ANOVA followed by Tukey post-hoc test was used for data analysis.

Next, the toxicity of 24HB was estimated under FBS-free conditions for adherently grown Y-79 cells, in accordance with the uptake experiments. At a 2 nM concentration, uncoated 24HB (NP0) as well as coated 24HB at coating ratios NP1, NP2 and NP3 were well-tolerated by Y-79 cells (Figure 4b). Free PEG5K-K10 (no 24HB) though, corresponding to the amounts used for coatings of NP2 and NP3, started to reduce the cell viability. Based on two-way ANOVA results, cell viability was both significantly affected by the presence of 24HB (*F*(1,16) = 45.924, p < 0.0001) and of PEG5K-K10 (*F*(3,16) = 11.410, p = 0.0003). The interaction between 24HB and PEG5K-K10 was also statistically significant (*F*(3,16) = 7.690, p = 0.002). According to the Tukey HSD post-hoc test, for NP1-coated 24HB, the means did not significantly differ between the coated DON and its equivalent amount of free polymer (101.9±11.2% vs. 99.8±7.5%, p = 1). Yet, for NP2-24HB and its equivalent amount of free polymer (107.8±9.8% vs. 81.7±5.6%, p = 0.019), and for NP3-24HB and its equivalent free amount of polymer (104.3±6.5% vs. 59.0±7.5%, p < 0.0001), they differed increasingly with higher concentration of free PEG5K-K10. Uncoated and coated 24HB at any tested NP ratio did not reveal any significant differences in viability to each other. Thus, the complexation of cationic polymers with DONs seemed to reduce their toxicity, as also previously reported.^91^ Even at high concentrations of 10 nM, 24HB and NP2-24HB were well tolerated in Y-79 cells (Table SI3), in line with previous reports.^25^ This indicated a strong electrostatic interaction between the oligolysine and the 24HB surface, preventing the detachment of the coating polymer in low ionic strength physiological conditions.^77^ As long as PEG5K-K10 was not freely available in solution, its toxic effects were curbed, underlining its suitability as a coating polymer for DONs.

### Single particle tracking in vitreous in ex vivo porcine eyes

Considering that the positively charged PEG5K-K10 polymer coating of 24HB DONs was well-tolerable and pivotal in enabling cell uptake, we next investigated whether the coating might also affect its diffusion in porcine vitreous. To this aim, uncoated 24HB-F and coated 24HB-F at coating ratios NP2 and NP5 were injected into enucleated porcine eyes, and the trajectories of the fluorescent particles were recorded to extract corresponding diffusion coefficients (Figure 5, Figure SI11). We occasionally observed a large discrepancy in the diffusion coefficients of different eyes that got injected with the same DONs and even encountered some mobility differences within the same particle population extracted from the same eye. This resulted most likely from the biological variability between eyes as well as from the disturbance of the vitreal structure integrity during manual preparation and injection of the pig eyes. Furthermore, only particles which happened to be on the imaging focal plane could be recorded, leading to only a fraction of injected DONs being available for analysis.

**Figure 5.**
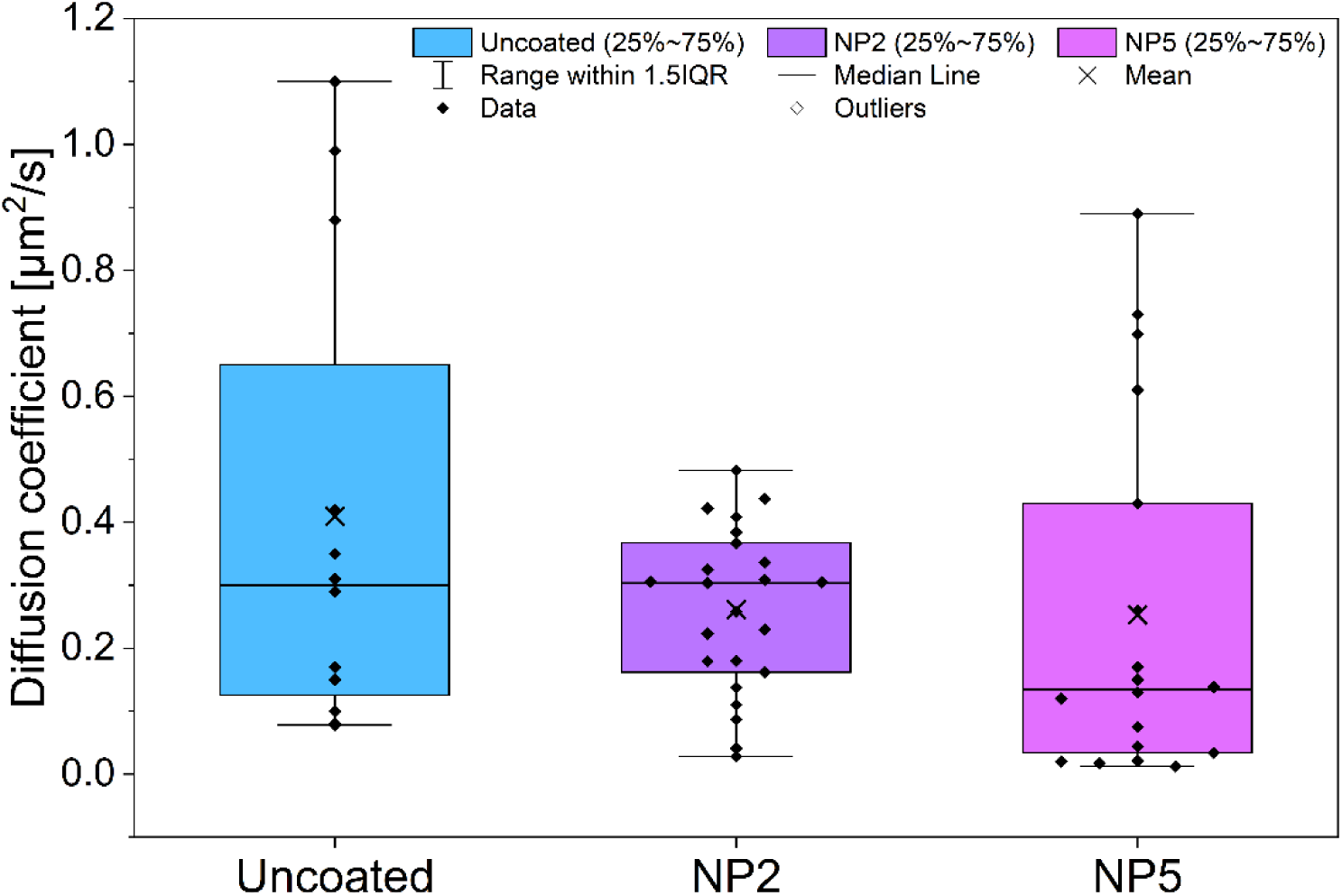
Distribution of diffusion coefficients for uncoated and coated 24HB-F at coating ratio NP2 and NP5, extracted from single particle tracking in vitreous of ex vivo porcine eyes. Four to five eyes for each DON type were examined, and all diffusion coefficients extracted were displayed.

Generally, the interquartile range of the diffusion coefficients values (encompassed by the box) fell for all DONs into the same range, indicating that they moved through vitreous similarly. For uncoated 24HB-F, the diffusion coefficients were shifted to slightly higher values (0.13-0.65 µm^2^ s^-1^) compared to NP2-24HB-F (0.16-0.37 µm^2^ s^-1^) and NP5-24HB-F (0.03-0.43 µm^2^ s^-1^). When comparing medians, however, the median diffusion coefficient of 0.13 µm^2^ s^-1^ for NP5-24HB-F was lower than for uncoated or NP2-24HB-F (both 0.30 µm^2^ s^-1^). Also, for NP5-24HB-F, several measurements indicated diffusion coefficients below 0.05 µm^2^ s^-1^, corresponding to some DONs being stuck in the vitreous while a fraction remained mobile. This matches the results by Käsdorf *et al.*, who reported that the penetration and diffusion of molecules and nanoparticles in bovine vitreous was dependent on their charge and that there was always a restricted and a mobile fraction observable, where the restricted fraction increased with positive charge.^58^ While the wide spread of diffusion coefficients made it difficult to draw clear conclusions, the positive charge from the PEG5K-K10 coatings would appear to not completely hinder the diffusion of DONs in porcine vitreous. Furthermore, we verified that despite the polyanionic nature of vitreal components such as hyaluronic acid and heparan sulfate, the coatings remained attached to DONs and could stabilize DONs in vitreous (Figure SI12). This is in line with observations of electrostatically assembled polyplexes that remain intact in vitreous as well.^92^ Tentatively, these results suggested that higher coating ratios such as NP5 that would most likely correspond with higher apparent cationic surface charge, might reduce nanoparticle mobility through the polyanionic, gel-like matrix of the vitreous, resulting in coated DONs getting immobilized more easily. On the other hand, lower coating ratios such as NP2 did not display largely different mobility behavior compared to uncoated 24HB, indicating that they can traverse the vitreous barrier.

This aligns well with reports from Tavakoli *et al.*, demonstrating that vitreal mobility of lipid nanoparticles is size- and charge-dependent, favoring anionic and neutral nanoparticles.^59^ In contrast, large neutral (>200 nm in diameter) and cationic lipid nanoparticles are restricted in their diffusion or immobilized,^59^ though PEGylation restores some mobility and reduces particle aggregation.^57,59,92–94^ Non-PEGylated, anionic liposomes in the ∼120-140 nm size range appeared slightly more mobile (0.37-0.6 µm^2^ s^-1^) compared to our uncoated 24HB,^59^ possibly due to shape differences, though tubular shapes have been suggested to be effective for transversing the vitreal gel, too.^57,95^ However, our coated 24HB DONs seemed less immobilized compared to PEGylated cationic liposomes (∼110-135 nm, 0.014-0.042 µm^2^ s^-1^).^59^ Perhaps, the PEG5K units of the coating reduced the direct interaction with the glycosaminoglycans and promoted DONs’ mobility more than the PEG2K modification for cationic liposomes. Alternatively, the liposomes were just more positively charged than our coated 24HB DONs. There seems to be some threshold for positive charge that would affect the degree of diffusion restriction for cationic compounds and particles,^58,96^ but perhaps also other factors like protein corona formation, particle architecture, and specific nanoparticle-vitreous interactions could play an important role.^59,97^ This would explain other similar accounts of positively charged nanoparticles where less dramatic effects of the zeta-potential on particle diffusion was observed.^96–98^

Generally, other kinds of nanoparticles have demonstrated higher mobility in similar experimental set-up, though direct comparison of diffusion coefficients warrants some caution.^59,97,98^ Investigating smaller or perhaps more compactly structured DONs could enhance their vitreal mobility, since anionic and neutral lipid nanoparticles under 50 nm size were barely restricted. Considering that many prevalent eye diseases affect elderly people, and vitreous humor undergoes progressive liquefaction and aggregation of collagen fibrils during the aging process,^99^ the mobility of nanoparticles might likely even increase, brought forth not only by vitreal fluid convection but by higher diffusivity.^59,100^ Ultimately, particle tracking in ex vivo porcine eyes gives only an approximation and does not necessarily directly translate to human vitreous,^101^ though at least the charge-selective properties of vitreous are preserved across different species.^58^ Besides vitreal diffusion, intravitreally injected nanoparticles also need to show sufficient ocular retention in vivo and overcome other barrier structures such as the inner-limiting membrane to reach retinal target tissue, often a limit for positively charged nanoparticles like polyplexes and those larger than 10-25 nm.^54,102–104^ Hence, further investigations and optimizations of DONs in regards to other ocular barrier structures and in vivo application is necessary in the future. In conclusion from this current work, PEGylated coatings for DONs seem compatible for their mobility in vitreous and settling for a coating ratio of NP2 or higher, which drives cell uptake, and below NP5 to curb the loss of particle mobility, could improve the odds of overcoming the vitreous barrier, balancing both properties of coated 24HB DONs for ocular application.

## Conclusion

Ocular diseases like retinoblastoma and other highly prevalent conditions like age-related macular degeneration and diabetic retinopathy threaten the patient’s eyesight or even life,^50–53,61^ and thus require effective delivery and treatment strategies. DONs are promising nanocarriers for ocular drug delivery applications as they offer precise and predictable tailoring opportunities for overcoming ocular barrier structures. In this work, we demonstrated that the limited cell uptake and structural stability of 24HB DONs in cell media and vitreous could be improved by an electrostatically assembled, commercial polymer coating. Thereby, Y-79 cells exhibited a distinct charge preference, where cell internalization of coated DONs was only observable at NP ratio ∼2 or higher. Furthermore, longer incubation and higher coating ratios increased the cell association of coated DONs. PL3 peptides were investigated as promising targeting ligands with DONs, as the target NRP-1 receptors are not only expressed in ocular cells^75^ but also widely in cancer cells.^63,64^ However, no additional benefit of the targeting moiety for cell uptake was observed in this study, possibly due to the necessity and impact of the coating driving the cell association, or due to the lack of proper receptor binding. We have discussed several considerations, and further investigations will offer the opportunity to differentiate between different compounding factors and to improve the DONs particle design and set-up. Generally, PEG5K-K10-coated DONs were mostly well-tolerable in ocular cell lines. Single particle tracking of DONs in ex vivo porcine eyes demonstrated their diffusive mobility in the polyanionic environment of the vitreous, even upon polymeric coating. This suggested that the charge-dependent trapping of cationic nanoparticles in the vitreous was not that strongly pronounced for PEG5K-K10-coated DONs in the coating ratio range that would seem suitable for cell uptake enhancement, though coating ratios below NP5 would be favorable. Furthermore, the coated DONs remained coated and stable in vitreous. These are important properties of DONs for ocular delivery, since intravitreal injection – one of the most common administration routes for targeting the posterior segment of the eye – leads to immediate exposure of the nanoparticles to the vitreous humor. Further studies are necessary to examine and modify the properties of DONs to overcome also other intraocular barrier structures like the inner limiting membrane and reaching their target tissue. Given their customizability and compatibility with a plethora of therapeutic and targeting molecules, DONs offer a great tailorable platform for systematic investigation and development of effective ocular delivery systems in the future.

## Experimental Section

### Preparation and characterization of 24-helix bundle (24HB) DNA origami nanostructures (DONs)

Using a thermal ramp, 24HB DONs were assembled in a one-pot reaction in 1×folding buffer (1×FOB, comprising of 1×Tris-acetate-EDTA buffer (1×TAE, containing 40 mM Tris, 20 mM acetic acid, 1 mM EDTA) and 17.5 mM MgCl_2_), as previously reported (details in Supporting information (SI), Section SI1).^76^ To create fluorescently labelled structures, selected DNA staple strands located at the ends of the DONs were exchanged with elongated strands, displaying a 22-nucleotide long overhang sequence. Those ssDNA overhangs were annealed in a second step, after folding and purification of the DONs, with sequence-complementary Atto488-modified DNA strands (Integrated DNA Technologies, USA, Section SI1).^60^ Similarly for peptide-carrying structures, selected staple strands were replaced with staple strands bearing a specific overhang-sequence to enable post-assembly functionalization with custom peptide PL3-modified DNA strands (Eurogentec, Belgium, CAGRGRLVR (PL3 peptide) linked *via* N-terminal cysteine and SMCC-linker to 5’ Amino-C6-TCC ACC ACA CCA CCA CCA CAA AA-3’, Section SI1). In detail, after preparation of the respective DONs, excess DNA staple strands were removed *via* polyethylene glycol (PEG) precipitation,^105^ and DONs were redissolved 1×FOB on a shaker (600 rpm, 20 °C). Subsequently if needed, DONs were mixed and annealed with a 10× excess of the functionalized strands (per available attachment site): 24HB-F DONs, allowing for the attachment of 12 Atto488-modified DNA strands on each end (24 in total), were prepared by cooling the annealing reaction in 1×FOB from 40 °C to 20 °C (- 0.1 °C/40 s),^60^ while 24HB-RFLP DONs with 12 positions for Atto488-modified strands on one end (RF) and 12 positions for PL3-modified strands (LP) on the other end, were kept overnight at room temperature. Similarly, 24HB-RF with attachment strands for LP (peptide-free control of 24HB-RFLP) was prepared. Respectively, excess DNA strands with attached fluorophore or peptide were removed *via* PEG precipitation, and 24HB-F resuspended in 1×FOB on a shaker at 20 °C, 24HB-RFLP and 24HB-RF statically at 4 °C. Fluorescent DONs were shielded from direct light during storage and experiments.

Proper folding, attachment of fluorophore and/or peptide, stability and removal of excess DNA strands were verified *via* agarose gel electrophoresis (AGE) and transmission electron microscopy (TEM, Section SI1). For AGE, samples were mixed with 6× loading dye solution (Sigma Aldrich, Germany), loaded onto a 2% (w/V) agarose gel with 0.46 µg mL^-1^ ethidium bromide, run in running buffer (1×TAE, 11 mM MgCl_2_) for 50 min at 90 V on ice and imaged using iBright FL1500 Imaging Systems (Invitrogen, USA). For TEM, samples were deposited on glow-discharged Formvar carbon-coated copper grids (FCF-400-CU, Electron Microscopy Sciences, USA), negatively stained with 2% (w/V) uranyl formate solution (pH-adjusted with 25 mM NaoH) and imaged on Hitachi HT7800 Transmission Electron microscope (100 kV acceleration voltage, Hitachi, Japan). AGE and TEM images were selected and cropped as needed. Concentration *c* (in mol L^-1^) of 24HB DONs was estimated based on its absorbance at 260 nm, using Lambert-Beer law (A= ε*cl*, *l*= 1 cm for NanoDrop One (Thermo Scientific, USA)). Considering the number of hybridized and non-hybridized nucleotides, the molar extinction coefficient ε was approximated for 24HB (1.05×10^8^ M cm^-1^), for 24HB with free overhangs (intermediate DON for 24HB-F and 24HB-RFLP, 1.083×10^8^ M cm^-1^), for 24HB-RF with free overhangs for LP (1.093×10^8^ M cm^-1^), and for 24HB-F and 24HB-RFLP (1.104×10^8^ M cm^-1^).^106^

### Preparation of coated 24HB DONs and removal of coating

To assemble a positively charged coating onto negatively charged 24HB DONs,^25^ we purchased methoxy-poly(ethylene glycol)-block-poly(L-lysine hydrochloride) (mPEG_5K_-b-PLKC_10_, Lot: 050-KC010-109) from Alamanda Polymers (USA), in the following referred to as PEG5K-K10, and stored it at -20 °C under argon. According to the Certificate of Analysis, the number average molecular weight by NMR was 6800 Da, the PDI from gel permeation chromatography 1.075. 10 mM aliquots of PEG5K-K10 in Milli-Q water were stored at -20 °C and sonicated for at least 2 min before use. Coating reactions were prepared in 1×FOB by mixing DONs with the PEG5K-K10 solution at the desired final concentration and N-to-P ratio (NP, charge ratio reflecting the number of positive charges from the protonated amine groups (N) in the lysines compared to the number of negative charges from the phosphate groups (P) in DNA, see Section SI2). They were rested at room temperature for at least 30 min, before further use or storage at 4 °C. Note, coated 24HB-RFLP and its peptide-free control 24HB-RF were coated both at a given NP ratio for comparability with the same quantities of polymer (as calculated for 24HB-RFLP), despite 24HB-RF having overall less negative charges, since no PL3-DNA strands were hybridized.

To remove the PEG5K-K10 coating from DONs, heparin sodium salt from porcine intestinal mucosa (Sigma Aldrich, Germany, 2 mM stock solution prepared in Milli-Q water) was added to coated DONs in excess (usually 50× in relation to the positive charges) and incubated at room temperature for at least one hour, unless mentioned otherwise. Coated DONs and the removal of the coating were visualized on a 2% agarose gel.

### Cell culture

The retinoblastoma cells Y-79 were a kind gift from Mika Reinisalo at the School of Pharmacy, University of Eastern Finland and originally purchased from the European Collection of Authenticated Cell Cultures. Y-79 cells were grown in suspension in TC-treated T75-flasks (Sarstedt, Germany) at 37 °C with 5% CO_2_, split twice per week and maintained between passage 16 and 48. Cell media (CM) consisted of RPMI-1640 (Gibco, USA, 21875-034) and was supplemented (CM_suppl_) with 10% fetal bovine serum (FBS, Gibco, USA), 1% glutamine (200 mM, Gibco, USA) and 1% penicillin-streptomycine (PS, 5 000 IU mL^-1^, Gibco, USA). Mycoplasma tests were performed when necessary. For most experiments, Y-79 cells were seeded in FBS-containing CM_suppl_ and grown adherently, then treated with DONs under FBS-free conditions to minimize background signal, enzymatic degradation of DONs and the detachment of the attached Atto488 fluorophores and peptides. To grow Y-79 cells adherently, the surface of the cell culture plates was pre-treated with 1 mg mL^-1^ poly-L-lysine (PLL) solution (30 000-70 000 MW poly-L-lysine hydrobromide, Sigma Aldrich, Germany, P2636) in borate buffer (pH 8.5) for at least 30 min at RT and washed twice with water before seeding cells.

### Uptake studies with flow cytometry

TC-treated 12-well plates (Sarstedt, Germany) were pre-treated with 460 µL/well of 1 mg mL^-1^ PLL solution for at least 30 min at RT and washed with 2 mL of water, twice. 220 000 Y-79 cells were seeded in 1 mL/well and let attach overnight at 37 °C with 5% CO_2_. The following day, the FBS-containing CM_suppl_ was exchanged with CM 1% PS (RPMI-1640, 1% PS), by removing 800 µL of CM and replacing it with 800 µL of CM 1% PS, twice. 250 µL of DON treatments in 1×FOB was spiked into the wells at a final DON concentration of 1.5 nM at a 20% (V/V) proportion and incubated at 37 °C with 5% CO_2_ for 24 h. To recover the cells for flow cytometry analysis, treatments were removed, cells washed with 2 mL of 1×Dulbecco’s phosphate-buffered saline (DPBS, Gibco, USA) and detached with 0.3 mL of 1×TrypLE Express Enzyme (Gibco, USA, 12604-021) for 7-10 min at 37 °C, before collecting the cell suspension in 1 mL of CM_suppl_ in 1.5 mL Eppendorf tubes. The cells were pelleted and washed twice, by centrifuging at 150 g for 4 min, aspirating the supernatant and resuspending the cell pellet in 1×DPBS. The cells were recovered after a third centrifugation directly in 0.5 mL of eBioscience Flow Cytometry Staining Buffer (Invitrogen, USA, 00-4222-26), filtered through the cell-strainer cap (mesh size: 35 µm) into Falcon 5 mL polystyrene round-bottom tubes (Corning, USA, 352235) and stored at 4 °C till measurement. Measurements were performed on the BD LSRFortessa Analyser (BD, USA) using the BD FACSDiva Software v 8.0. The flow cytometer was calibrated daily with CST beads before starting measurements, the blue laser (488 nm) was used with the detector wavelength 530/30 (BP filter: 515-545 nm, preceding LP filter 505 LP). Based on the buffer-control sample, voltages, gates and FSC area scaling were adjusted daily and set to include around 20 000 live cells. The analysis was performed on FlowJo v.10.10.0 software (BD, USA) by first gating the live cells by size and granularity (FSC-A vs. SSC-A), selecting the single cell population in the doublet-discrimination plot (FSC-A vs. FSC-H) and observing the fluorescence signal in the Atto488 channel (example of gating strategy shown in Figure SI13). The fluorescence overlap of the gate that was to include cells positive for Atto488 was set to max. 0.5% for the buffer-control sample. Experiments were usually performed with 1-2 technical replicates on 2-3 separate days (n) depending on the experiment.

### Live confocal imaging for uptake studies

The wells of the CellView cell culture slide (TC, glass bottom, Greiner Bio-One, Austria, 543078) were pre-treated with 1 mg mL^-1^ PLL solution (45 µL/well) for at least 30 min at RT and washed twice with 200 µL of water, before seeding the cells. 15 000 Y-79 cells were added in 100 µL/well and let attach overnight at 37 °C with 5% CO_2_. The next day, the FBS-containing CM_suppl_ was replaced with FBS-free CM 1% PS (RPMI-1640, 1% PS) by replacing 80 µL of CM with CM 1% PS, twice. Atto488-labelled DON treatments in 1×FOB were spiked to the cells to a final concentration of 10 nM at a 20% (V/V) proportion and incubated at 37 °C with 5% CO_2_ for 24 h. LysoTracker Deep Red (Invitrogen, USA) was mixed into respective wells (final concentration in well: 100 nM) for the last hour of the incubation. The localization of fluorescent DONs in Y-79 cells was then visualized with an inverted confocal laser scanning microscope (Leica TCS SP8 STED 3X CW 3D STED (Stimulated Emission Depletion), Germany) using a 63×/1.20 (water, wd 0.3 mm) objective. Atto488 (λ_exc_= 502 nm, λ_em_= 522 nm) was excited with an argon 488 nm laser, LysoTracker Deep Red (λ_exc_= 647 nm, λ_em_= 668 nm) with a HeNe 633 nm laser. The final images were exported and analyzed using LAS X software (version 3.7.4.23463).

### Cell viability assessment with alamarBlue Assay

The toxicity of free PEG5K-K10 (final concentrations: 0.5-100 µM) was tested in FBS-containing CM_suppl_ on Y-79 cells in suspension, while PEG5K-K10-coated 24HB (final DON concentration: 2 nM or 10 nM) and corresponding controls were studied under FBS-free conditions on adherently grown Y-79 cells. In both cases, 20 000 Y-79 cells were added in 100 µL/well of FBS-containing CM_suppl_ on a 96-well plate and grown overnight. Y-79 cells were attached if needed by pre-treating the wells with 45 µL of 1 mg mL^-1^ PLL solution for at least 30 min at RT and washing with water (200 µL/well) twice prior cell seeding. The following day, the CM_suppl_ was replaced for adherently grown cells with FBS-free media CM 1% PS (RPMI-1640, 1% PS) by gently removing 80 µL of CM and replacing it with 80 µL CM 1% PS, twice. Then, both suspension and adherent cells were treated respectively by adding 25 µL of PEG5K-K10 or coated 24HB in 1×FOB to the cells at a final proportion of 20%(V/V) and incubated at 37 °C with 5% CO_2_ for 24 h. The next day, alamarBlue HS Cell Viability Reagent (Invitrogen, USA, A50100) was prepared as 6×solution in CM 1% PS and spiked into the wells (to a final concentration: 1×) and incubated at 37 °C with 5% CO_2_ for 4 h. To read-out fluorescence, 100 µL of each well was transferred after thorough mixing into a black-walled well plate (Thermo Scientific, USA, 265301) and measured with Varioskan LUX (Thermo Scientific, USA, top optics, λ_exc_= 560 nm, λ_em_= 590 nm). The experiment was repeated on three separate days (n=3) with each treatment condition in a technical triplicate. The cell viability was expressed as % viability relative to the buffer-treated control.

### Single particle tracking in porcine vitreous

As in the previously reported protocol,^59^ pig eyes acquired from the abattoir (HKScan Finland Oyj, Finland) were cleaned from excess tissue, rinsed with 70% ethanol and stored in 1×PBS at 4 °C. To inject an eye, the anterior part of the eye, including the lens was excised, exposing the vitreous humour. Then, 50-100 µL of a 30 nM DONs solution (uncoated or coated 24HB-F) in 1×FOB was injected *via* a 30 G insulin syringe (BD, USA) into the vitreous in the center of the eye (ca. 0.5 cm deep) and briefly rested. The eye was flipped onto a 35 mm microwell dish with 14 mm diameter glass window (MatTek Corporation, USA) and fixed to the dish with cyanoacrylate glue (Henkel Corp., USA). The imaging was performed at room temperature with a spinning disc confocal microscope. Two imaging systems were used to record time series with 50 ms resolution. Microscope Zeiss Axio Observer Z1 was equipped with Zeiss 63×/1.2 W C-Apochromat objective. In case of Marianas (Intelligent Imaging Innovation (3i), USA), the 488 nm/ 150 mW solid state laser and Yokogawa CSU-X1 M1 5000 rpm spinning disk were used as a light source. In case of Aurox Clarity (Aurox Ltd., UK), LED light source CoolLED pE-400 max along with FITC CC00302 filter were used. The trajectories were analyzed with Imaris v. 10.1 software (Bitplane AG, Switzerland) and the particle mean square displacement (MSD) was computed using the @msdanalyzer MATLAB plug-in (MathWorks Inc., USA).

The diffusion coefficient of DONs in vitreous (D) was calculated using the equation D=*α/(*2*d)*, where *α* is a slope of mean square displacement vs. time dependence and *d* is 2 for two-dimensional diffusion.

### Statistical methods

Statistical analysis was performed with Origin 2026. The data is presented as mean with standard deviation and where applicable, was analyzed using 2-way ANOVA followed by Tukey post-hoc test.

## Acknowledgements

We acknowledge financial support through the European Research Council Consolidator Grant (PADRE, 101001016), the Research Council of Finland (flagship GeneCellNano, project no. 361647, project no. 355520), Jane and Aatos Erkko Foundation, Silmäsäätiöiden tohtoritutkijapooli, the Estonian Research Council grant PRG3076, Mobilitas 3.0 Program under the framework of the JPIAMR - Joint Programming Initiative on Antimicrobial Resistance (MOB3ERA1), and ERA Chair MATTER from the European Union’s Horizon 2020 research and innovation program under grant agreement No 856705. We also thank Viikki Flow Cytometry unit, Light Microscopy Unit, and Electron Microscopy Unit at the University of Helsinki, supported by HiLIFE and Biocenter Finland, for access to their facilities and support during data acquisition and analysis. A.K. would like to thank the Biohybrid Materials Group under Prof. Mauri Kostiainen at Aalto University for their support and access to their facilities. Graphical abstract and Figures 1a and 1d were assembled with BioRender.com.

## Conflicts of Interest

The authors declare no conflicts of interest.

## Data Availability Statement

The data that support the findings of this study are available from the corresponding author upon reasonable request.

## Graphical Abstract

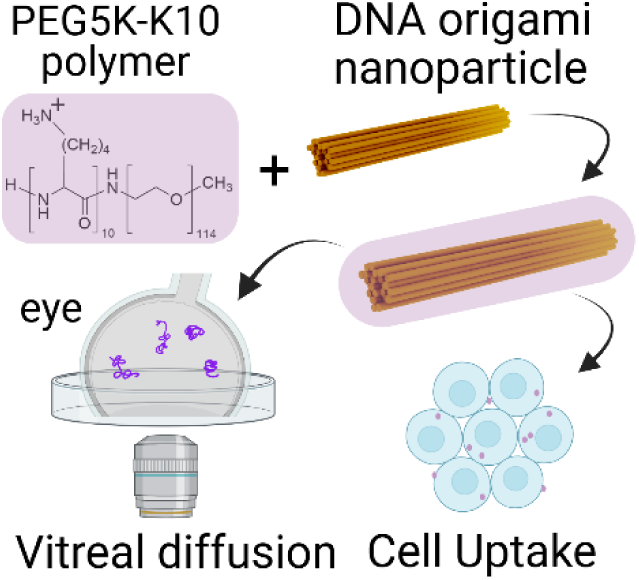

### Table of Contents

We show that polymeric coating can enable DNA origami nanoparticle internalization into ocular retinoblastoma cells. Not only are the coated nanoparticles well-tolerable to the cells, but they also remain mobile in the vitreous of a porcine ex vivo model. Thus, they could offer a promising drug delivery platform for treating ocular diseases in the future.

## Notes

### Competing Interest Statement

The authors have declared no competing interest.

