## Supporting Information for "DNA origami uptake in Y-79 retinoblastoma cells driven by oligolysine coating"

A. Klose, Z. Gounani, S. Raik, A. Koivuniemi, T. Laaksonen
Division of Pharmaceutical Biosciences, Faculty of Pharmacy, University of Helsinki, Helsinki, Finland

S. Korhonen, M. Reinisalo, T. Lajunen
School of Pharmacy, Faculty of Health Sciences, University of Eastern Finland, Kuopio, Finland

S. Korhonen
Laboratory of DDS Design and Drug Disposition, Graduate School of Pharmaceutical Sciences, Tohoku University, Sendai, Japan

V. Linko
Institute of Technology, University of Tartu, Tartu, Estonia

T. Laaksonen
Chemistry and Advanced Materials, Faculty of Engineering and Natural Sciences, Tampere University, Tampere, Finland

E-mail addresses of corresponding authors:

### SI1. Preparation and characterization of 24HB DNA origami nanostructures (DONs)

One-pot folding reactions for 24HB DONs were prepared in 1× folding buffer (1×FOB), consisting of 1×Tris-acetate-EDTA buffer (1×TAE: 40 mM Tris, 20 mM acetic acid, 1 mM EDTA) and 17.5 mM $MgCl_2$.[1] To this aim, scaffold strands p7560 (Tilibit, Germany) at a final concentration of 20 nM were mixed with ~10× excess of staple strands (Integrated DNA Technologies, USA) and annealed in the GeneAmpPCR System 2700 (Applied Biosystems, USA) by controlled heating (30 s at 65 °C) and subsequent controlled cooling (from 65 °C to 59 °C at a rate of -1 °C/15 min, from 59 °C to 40 °C at a rate of -0.1 °C/18 min, stored at 20 °C till recovery).

Ijäs *et al.* previously reported the design and staple strand sequences for 24HB.[1] For 24HB DONs that were decorated with fluorophores (Atto488) and/or peptides (PL3), specific staple strands located at the ends of the rod-like 24HB structure were replaced with staple strands of extended length and sequence, that could specifically hybridize with Atto488-modified or PL3-modified oligonucleotides. Locations of the exchanged staple strands[2] and their sequences for Atto488 attachment have been reported previously.[3] Relevant sequences can be found in **Table SI1**.

The thymine next to the fluorophore or peptide attachment was left unpaired, enabling a 22 base pair-binding (<u>underlined</u>) with the overhanging staple strands from 24HB.

**Atto488-modified oligonucleotide** (23 nucleotides long):

/5ATTO488N/T<u>GG GAG AGG AGG AGG AGA AAA AA</u>

**PL3-modified oligonucleotide** (23 nucleotides long):

/5peptide/T<u>CC ACC ACA CCA CCA CCA CAA AA</u>

The PL3 peptide is linked *via* an added N-terminal cysteine (**C**AGRGRLVR) and SMCC-linker to 5' Amino-C6- TCC ACC ACA CCA CCA CCA CAA AA-3'. PL3-modified DNA strands were custom-ordered (Eurogentech, Belgium), dissolved in Milli-Q water and stored in small aliquots at -20 °C or -80 °C.

**Table SI1. Extended staple strand sequences of 24HB for the hybridization with Atto488-oligonucleotide or PL3-oligonucleotide.** The sequence complementary to the scaffold is shown in capital letters, while lowercase letters indicate the overhanging sequences. The overhang responsible for hybridizing with the Atto488- or PL3-modified oligonucleotide is italicized. For 24HB-F, r1-r23 (highlighted in blue) and l0-l22 (highlighted in green) were incorporated into the 24HB structure. For 24HB-RFLP, r1-r23 (highlighted in blue) and l0p-l22p (highlighted in red) were selected. (Continued on the next page)

(Continuation: Table SI1)

| # | **Sequence** |
|---|---|
| For hybridizing with Atto488-modified oligonucleotides | |
| r1 | ttttttttGCAAGGATAAAACAATTCTGC*ttttttctcctcctcctctccc* |
| r3 | ttttttttAAGCTAAATCGGAATAACCTG*ttttttctcctcctcctctccc* |
| r5 | ttttttAGCATTAACATCCAATTTCTACTAATAGTAGT*ttttttctcctcctcctctccc* |
| r7 | ttttttttTCATTGCCTCCTCAGAGCATA*ttttttctcctcctcctctccc* |
| r9 | ttttttttATAAATTAACTTTATTTCAAC*ttttttctcctcctcctctccc* |
| r11 | ttttttAAGGGTGAGAAAGGCCGTAGGTAAAGATTCAA*ttttttctcctcctcctctccc* |
| r13 | ttttttttAGGTCACGTTGGTTCTAGCTG*ttttttctcctcctcctctccc* |
| r15 | tttttttttTTAAATGTGAGCGCTATCAGG*ttttttctcctcctcctctccc* |
| r17 | ttttttTCATTTTTTAACCAATTTTTTGTTAAATCAGC*ttttttctcctcctcctctccc* |
| r19 | ttttttAAATTGTAAACGTTAAGTATAAGCAAATATTT*ttttttctcctcctcctctccc* |
| r21 | GAAGATTTATTTTGCATTAAAAGGAACGTAGCCAGCTTTCATCAACA*ttttttctcctcctcctctcc c* |
| r23 | TACAAAGGAGTAACGGATTGACCGTAATGGGAT*ttttttctcctcctcctctccc* |
| l0 | CCAACGCTCCCTTAAAGAGTCCACTATTAAAGA*ttttttctcctcctcctctccc* |
| l2 | tttttttCCGCCTGGCCCTCTGTTTGATGGTGGTTCCG*ttttttctcctcctcctctccc* |
| l4 | ttttttttGCGAACTGATAGGATTGCCCTTCA*ttttttctcctcctcctctccc* |
| l6 | ttttttttAAATCGGCAAAAGCGGGGAGA*ttttttctcctcctcctctccc* |
| l8 | ttttttttACGTGGACTCCATTAATTGCG*ttttttctcctcctcctctccc* |
| l10 | AGCCAGGGTGGATGTTAAGCTTTACCGAGCTCACAATTCCAC*ttttttctcctcctcctctccc* |
| l12 | ttttttttTTGCGCTCAGATAAAGACGGA*ttttttctcctcctcctctccc* |
| l14 | ttttttttGGCGGTTTGGCATTTCACATA*ttttttctcctcctcctctccc* |
| l16 | CAGTGCCCTTCTAATCCTTAGCCAAAATGGAGTGACTCTATGATACC*ttttttctcctcctcctctcc c* |
| l18 | CTGCCATGGCTATTAGTCTTTAATGC*ttttttctcctcctcctctccc* |
| l20 | ttttttttAATCATTTCTCCTTGTCAACC*ttttttctcctcctcctctccc* |
| l22 | ttttttttGGATCCCCGGGTCTCAGGAGA*ttttttctcctcctcctctccc* |
| For hybridizing with PL3-modified oligonucleotides | |
| l0p | CCAACGCTCCCTTAAAGAGTCCACTATTAAAGA*ttttgtggtggtggtgtggtgg* |
| l2p | tttttttCCGCCTGGCCCTCTGTTTGATGGTGGTTCCG*ttttgtggtggtggtgtggtgg* |
| l4p | ttttttttGCGAACTGATAGGATTGCCCTTCA*ttttgtggtggtggtgtggtgg* |
| l6p | ttttttttAAATCGGCAAAAGCGGGGAGA*ttttgtggtggtggtgtggtgg* |
| l8p | ttttttttACGTGGACTCCATTAATTGCG*ttttgtggtggtggtgtggtgg* |
| l10p | AGCCAGGGTGGATGTTAAGCTTTACCGAGCTCACAATTCCAC*ttttgtggtggtggtgtggtgg* |
| l12p | ttttttttTTGCGCTCAGATAAAGACGGA*ttttgtggtggtggtgtggtgg* |
| l14p | ttttttttGGCGGTTTGGCATTTCACATA*ttttgtggtggtggtgtggtgg* |
| l16p | CAGTGCCCTTCTAATCCTTAGCCAAAATGGAGTGACTCTATGATACC*ttttgtggtggtggtgt ggtgg* |
| l18p | CTGCCATGGCTATTAGTCTTTAATGC*ttttgtggtggtggtgtggtgg* |
| l20p | ttttttttAATCATTTCTCCTTGTCAACC*ttttgtggtggtggtgtggtgg* |
| l22p | ttttttttGGATCCCCGGGTCTCAGGAGA*ttttgtggtggtggtgtggtgg* |

To remove excess DNA staple strands, the folding reaction was first diluted 1:3 (V/V) with 1×FOB, then 1:1 (V/V) with polyethylene glycol (PEG) precipitation buffer (1×TAE, 15% (w/V) PEG8000, 505 mM NaCl) before centrifuging at 14 000 g for 30 min.[4] After removing the supernatant, the DON pellet was redissolved in 1×FOB on a shaker at 20 °C at 600 rpm.

To prepare 24HB DONs with attached Atto488 fluorophores and/or peptide PL3, the respective purified 24HB base structure displaying the necessary attachment strands was mixed with Atto-modified and/or PL3-modified DNA strands at a 10× molar excess (per attachment site) and annealed. In case of 24HB-F, the annealing reaction was cooled from 40 °C to 20 °C (at a rate of -0.1 °C/40 s in T100 Thermal Cycler (Bio-Rad, USA) or GeneAmpPCR System 2700 (Applied Biosystems, USA)),[3] while 24HB-RF and 24HB-RFLP (when needed also 24HB-LP with attachment strands for RF) were left to anneal at room temperature overnight. Subsequently, DONs were purified from excess Atto488- or PL3-modified DNA strands *via* PEG precipitation (as described above) and redissolved in 1×FOB on a shaker at 20 °C at 600 rpm (24HB-F) or statically at 4 °C (24HB-RF, 24HB-RFLP).

The concentration (mol $L^{-1}$) of 24HB DONs was estimated according to Lambert-Beer law, as detailed in the main text.

To prepare samples for transmission electron microscopy (TEM),[5] formvar carbon-coated copper grids (FCF400-Cu, Electron Microscopy Sciences, USA) were treated at 25 mA for 45 s in the Emitech K100X Glow discharge unit. 3 µL of 4 nM DON samples were incubated on the grid for 4 min before blotting. For negative staining, the grids were subsequently flipped onto a 5 µL droplet of 2% (w/V) uranyl formate solution (UFO, pH-adjusted with 25 mM NaOH), immediately blotted, then flipped onto a 20 µL droplet of 2% UFO and incubated for 45 s, before final blotting and drying the grid. The grids were stored at RT in a grid box till imaging.

To check, the removal of excess DNA strands and incorporation of Atto488 and/or PL3, 24HB DONs were mixed with 6× loading dye solution (Sigma Aldrich), loaded onto a 2% agarose gel (containing 0.46 µg $mL^{-1}$ ethidium bromide) and run in 1×TAE, 11 mM $MgCl_2$ (running buffer) at 90 V for 50 min on ice. Gels were visualized using iBright FL1500 Imaging Systems (Invitrogen). For stability assessment, samples were incubated at 4 °C or 37 °C for 24 h. When needed, coated samples were decoated with heparin and subsequently run on an agarose gel (same conditions as above). Upon imaging, main bands were checked for changes and shifts as well as for loss of the Atto488 signal. (for details refer to the caption of the respective SI figures)

### SI2. Calculations for coating of PEG5K-K10-24HB and coating removal

Based on the number of nucleotides in the DON structure, we estimated for a regular 24HB DON (without any modifications) 15504 negative charges and for 24HB-F or 24HB-RFLP 16375 negative charges. The control DON 24HB-RF (with unoccupied attachment strands for LP) technically only carries 16099 negative charges but was coated as a control always with the same quantities as calculated for 24HB-RFLP.

According to the certificate of analysis, our PEG5K-K10 from Alamanda polymers (Lot: 050-KC010-109) had a number average molecular weight by NMR of 6800 and a degree of polymerization (number of monomeric units) of the poly-lysine block by NMR of 9. 10 mM PEG5K-K10 stock in MilliQ water were stored at -20 °C (prepared based on 6800 g/mol as molecular weight). The polymer was sonicated for at least 2 min before use and if needed, diluted further to a working stock.

#### Exemplary calculation for PEG5K-K10 amount needed for a specific NP coating ratio

E.g. preparation of coated 24HB-F at NP2 (20 µL, 3 nM)

1. Amount (n) of DONs:

$$n(DONs) = concentration\ of\ DONs * volume\ of\ coating\ reaction$$

$$n(DONs) = 3 * 10^{-9} \frac{mol}{L} * 20 * 10^{-6} L = 6 * 10^{-14}\ mol$$

2. Amount (n) of negative charges in the coating reaction:

$$n(negative\ charges) = n(DONs) * number\ of\ negative\ charges\ in\ one\ DON$$

$$n(negative\ charges) = 6 * 10^{-14}\ mol * 16375 = 9.825 * 10^{-10}\ mol$$

3. Amount (n) of positive charges required according to NP ratio:

$$n(positive\ charges) = n(negative\ charges) * NP\ ratio$$

$$n(positive\ charges) = 9.825 * 10^{-10}\ mol * 2 = 1.965 * 10^{-9}\ mol$$

4. Amount (n) of PEG5K-K10 required:

$$n(PEG5K - K10) = \frac{n(positive\ charges)}{number\ of\ positive\ charges\ in\ one\ PEG5K - K10\ molecule}$$

$$n(PEG5K - K10) = \frac{1.965 * 10^{-9}\ mol}{9} \approx 2.183 * 10^{-10}\ mol$$

5. Volume (V) of PEG5K-K10 stock required:

$$V\ (PEG5K - K10\ stock) = \frac{n(PEG5K - K10)}{concentration\ of\ PEG5K - K10\ stock}$$

$$V\ (PEG5K - K10\ stock) = \frac{2.183 * 10^{-10}\ mol}{100 * 10^{-6} \frac{mol}{L}} = 2.183 * 10^{-6}\ L \approx 2.18\ \mu L$$

The average number of negatively charged sulfate groups per heparin molecule was approximated to 71, assuming an average molecular weight of 18 000 g/mol for heparin and an average of 2.33 sulfate groups per each repeating IdoA(2S)-GlcNS(6S) disaccharide unit.[6] Heparin was prepared as 2 mM stock solution in MilliQ water and was stored at 4 °C.

Polyanionic heparin induces dissociation of the oligolysine-DON complex, resulting in the removal of the protective coating.

**Exemplary calculation for removal of PEG5K-K10 coating with heparin:**

E.g. Removal of coating from NP2-24HB-F (4 µL, 30 nM), heparin added at a 50× charge excess (compared to the coating).

1. Amount (n) of positive charges in the coating:
   (derived similarly as described in example above, here: $3.93*10^{-9}$ mol)
2. Amount (n) of negative charges required to decoat at a given charge excess:
$$n(negative\ charges) = n(positive\ charges) * charge\ excess$$
$$n(negative\ charges) = 3.93 * 10^{-9}\ mol * 50 = 1.965 * 10^{-7}\ mol$$
3. Amount (n) of heparin required:
$$n(heparin) = \frac{n(negative\ charges)}{number\ of\ sulfate\ groups\ per\ heparin\ molecule}$$
$$n(heparin) = \frac{1.965 * 10^{-7}\ mol}{71} \approx 2.768 * 10^{-9}\ mol$$
4. Volume (V) of heparin stock required:
$$V(heparin) = \frac{n(heparin)}{concentration\ of\ heparin\ stock}$$
$$V(heparin) = \frac{2.768 * 10^{-9}\ mol}{2 * 10^{-3}\frac{mol}{L}} = 1.384 * 10^{-6}\ L \approx 1.38\ \mu L$$

**SI3. TEM images of PEG5K-K10 coated 24HB DONs**

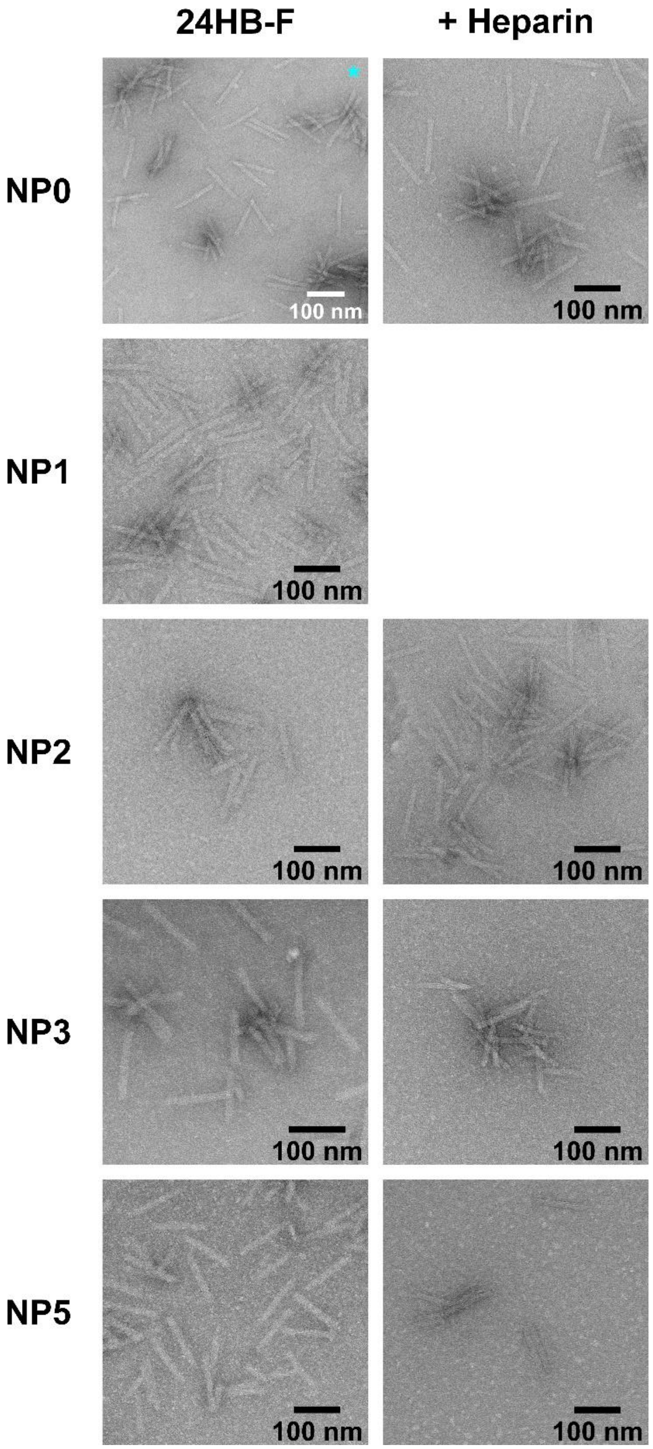

**Figure SI1. TEM images of 24HB-F coated at different NP ratios (left column) and after removal of the coating with 50× charge excess of heparin (right column).** Note that NP0 with cyan asterix shows 24HB-RFLP (instead of 24HB-F). NP0 + heparin was incubated with an excess of heparin equivalent for a NP5 coating. Majority of TEM images were cropped.

### SI4. Testing different amounts of heparin for removing PEG5K-K10 coating from DONs

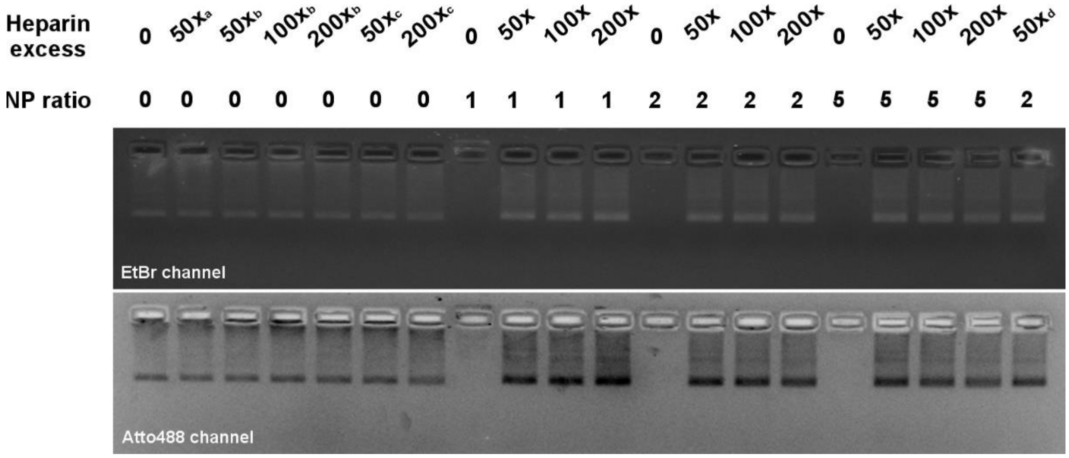

**Figure SI2. Screening of different Heparin amounts to remove PEG5K-K10 coating from 24HB-F at different NP coating ratios.** Heparin amounts were expressed as x-fold negative charge excess compared to the positive charges provided in the respective PEG5K-K10 coating (for more details refer to Section SI2). 20 µL of 3 nM coated 24HB-F were incubated with 8 µL of Heparin/water for an hour, before checking the samples on an agarose gel. For the uncoated controls (NP ratio 0), 24HB-F was mixed with different amounts of Heparin equivalent to the denoted charge excess given for NP1 (a), NP2 (b) or NP5 (c).The same bands of 24HB-F were observable both in the ethidium bromide (EtBr) and fluorescent channel (Atto488), indicating that 24HB-F remained stable in presence of the coating and the screened heparin amounts. Coated 24HB-F was immobilized in the gel pocket, while addition of heparin removed the polymer and led to the corresponding band on the gel. Complete removal of the PEG5K-K10 coating was achievable already at a 50x charge excess for all tested NP ratios. There appeared to be a minimal band shift after removal of the coating compared to the uncoated control that remained even at higher heparin concentrations, similar to a previous report.[7] Possibly, there was still some minimal interaction with the polymer in solution remaining. One hour of incubation was suitable for decoating DONs, but even shorter incubations as tested for the last sample (d, 25 min) seem feasible.

### SI5. Stability of 24HB DONs in cell media

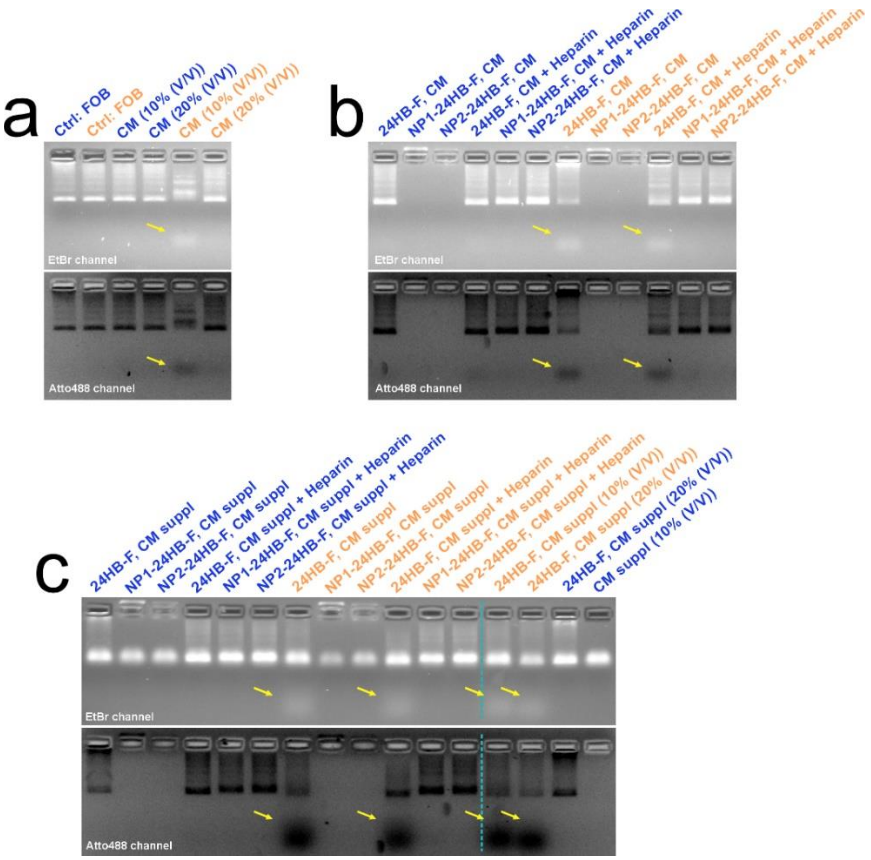

**Figure SI3. Stability of uncoated and PEG5K-K10 coated 24HB in cell media.** Reactions were incubated for 24 h at 4 °C (blue) or 37 °C (orange). **a.** Stability of uncoated 24HB-F was tested in 1×FOB (Ctrl) or in CM (RPMI-1640, 1% PS), with DONs in 1×FOB making up either 10% or 20% (V/V) (2 nM, 30 µL). 24HB-F appeared instable in CM at 10% (V/V) at 37 °C but stable at 20% (V/V). Most likely at 10% (V/V) dilution, the total Mg concentration in the reaction (2.11 mM) dropped below a critical threshold,[3] while at 20% (V/V) it was sufficient (3.82 mM). **b.** 24HB-F and coated 24HB-F at coating ratio NP1 and NP2 in 1×FOB were incubated in CM (RPMI-1640, 1% PS) at 10% (V/V) (3 nM, 20 µL). Coated samples were decoated with 50× charge excess of Heparin after the incubation, where indicated. Uncoated 24HB-F displayed loss of Atto488 attachments at 37 °C while coated DONs were stable. **c.** Uncoated 24HB-F and coated 24HB-F at NP1 and NP2 in 1×FOB were incubated in CM suppl (RPMI-1640, 10% FBS, 1% glutamine, 1% PS) at default 10% (V/V) or 20% (V/V) where indicated (3 nM, 20 µL). Coated samples were decoated in 30 min with 50× charge excess of Heparin after the incubation, where indicated. CM suppl gave an overlapping signal in EtBr channel where the band for 24HB-F was expected, so interpretation relied mainly on the Atto488 channel. In CM suppl at 37 °C, the loss of Atto488-tag was even more pronounced for all uncoated 24HB-F, regardless of the dilution proportion. Coated DONs remained stable, in line with previous reports that DNA extensions from DONs are well-protected and stabilized by PEG5K-K10 coatings.[8] For all samples, DONs were either diluted with water or Heparin after incubation, adjusted with $MgCl_2$ to equal total Mg amounts and diluted with 6×loading dye before running the gel (90 V, 50 min on ice). Gels were imaged in both the ethidium bromide (EtBr) and Atto488 channel.

### SI6. Uptake of coated 24HB DONs in Y-79 cells by flow cytometry measurements

**Table SI2. Percentage of cells positive for Atto488 from single Y-79 cell population during flow cytometry analysis.** Each treatment was studied on three separate days (repeat 1-3) in duplication. The readout of the duplicates is shown in the upper row, while the average of the duplicates of the same day is shown in bold underneath, respectively. Average and standard deviation (± s.d.) of the three repeats are presented in Figure 2bc of the main article.

<table>
<tr><th colspan="2"></th><th colspan="8">% of cells positive for Atto488</th></tr>
<tr><th>Incubation time</th><th>Sample</th><th colspan="2">Repeat1</th><th colspan="2">Repeat2</th><th colspan="2">Repeat3</th><th>Average</th><th>± s.d.</th></tr>
<tr><td rowspan="4">3 h</td><td rowspan="2">NP2-24HB-RF</td><td>9.98</td><td>13.4</td><td>5.28</td><td>7.09</td><td>51.7</td><td>56.6</td><td rowspan="2">24.01</td><td rowspan="2">26.25</td></tr>
<tr><td colspan="2">11.69</td><td colspan="2">6.19</td><td colspan="2">54.15</td></tr>
<tr><td rowspan="2">NP2-24HB-RFLP</td><td>19.6</td><td>22.8</td><td>8.57</td><td>13.1</td><td>59.6</td><td>62.2</td><td rowspan="2">30.98</td><td rowspan="2">26.43</td></tr>
<tr><td colspan="2">21.2</td><td colspan="2">10.84</td><td colspan="2">60.9</td></tr>
<tr><td rowspan="8">24 h</td><td rowspan="2">24HB-RF</td><td>8.95</td><td>9.28</td><td>8.04</td><td>10.7</td><td>17.3</td><td>13.9</td><td rowspan="2">11.36</td><td rowspan="2">3.67</td></tr>
<tr><td colspan="2">9.12</td><td colspan="2">9.37</td><td colspan="2">15.6</td></tr>
<tr><td rowspan="2">24HB-RFLP</td><td>1.48</td><td>1.5</td><td>1.71</td><td>1.67</td><td>1.23</td><td>1.26</td><td rowspan="2">1.48</td><td rowspan="2">0.22</td></tr>
<tr><td colspan="2">1.49</td><td colspan="2">1.69</td><td colspan="2">1.25</td></tr>
<tr><td rowspan="2">NP2-24HB-RF</td><td>49.3</td><td>45.2</td><td>43.8</td><td>45</td><td>81.6</td><td>79.1</td><td rowspan="2">57.33</td><td rowspan="2">19.98</td></tr>
<tr><td colspan="2">47.25</td><td colspan="2">44.4</td><td colspan="2">80.35</td></tr>
<tr><td rowspan="2">NP2-24HB-RFLP</td><td>49.9</td><td>48.4</td><td>48.3</td><td>47.4</td><td>87.2</td><td>86.4</td><td rowspan="2">61.27</td><td rowspan="2">22.12</td></tr>
<tr><td colspan="2">49.15</td><td colspan="2">47.85</td><td colspan="2">86.8</td></tr>
</table>

### SI7. ARPE-19 cell culture and exploratory uptake study with coated 24HB DONs by flow cytometry measurement

The ARPE-19 cells (human retinal pigment epithelial cells) were kindly gifted by Arto Urtti from the University of Eastern Finland and originally acquired from the American Type Culture Collection (ATCC). ARPE-19 cells were grown adherently in TC-treated T75 flasks (Sarstedt) at 37 °C with 7% or 5% $CO_2$ and passaged 1-2 times per week. The cell media (CM) consisted of DMEM/F12, Hepes (Gibco, 31330-038), 10% fetal bovine serum (FBS, Gibco) and 1% penicillin-streptomycine (PS, 5 000 IU $mL^{-1}$, Gibco). Mycoplasma tests were performed when needed.

#### Flow cytometry for uptake study (ARPE-19)

105 000 ARPE-19 cells were grown in TC-treated 12-well plates (Sarstedt) in 1 mL/well overnight at 37 °C with 5% $CO_2$. The following day, the FBS-containing CM was removed, the cells were washed with 1 mL of 1×Dulbecco's phosphate-buffered saline (DPBS, Gibco), DON treatments in 1×FOB at a 20% (V/V) proportion in CM 1%PS (DMEM/F12, 1% PS) were added and incubated at 37 °C with 5% $CO_2$ for 24 h (final DON concentration: 1 nM of 24HB-F for their higher number of fluorophores per DON). Then, the treatments were removed, the cells washed with 2 mL of 1×DPBS and detached with 0.3 mL of 1×TrypLE Express Enzyme (Gibco, USA, 12604-021) for 7 min at 37 °C. The cell suspension was collected in 1 mL CM in 1.5 mL Eppendorf tubes for washing. The cells were repeatedly centrifuged (150 g for 4 min) and washed twice, by removing the supernatant and mixing the cell pellet in 1×DPBS. After a third centrifugation, the cells were resuspended in 0.3-0.35 mL of eBioscience Flow Cytometry Staining Buffer (Invitrogen, 00-4222-26), filtered through the cell-strainer cap (mesh size: 35 µm) into 5 mL polystyrene round-bottom tubes (Falcon, 352235) and stored at 4 °C till measurement. The flow cytometry measurements and analysis were performed similarly as for Y-79 cells (refer to the main article's method section). For AREP-19, exploratory experiments were performed only once (n=1) in technical triplicate.

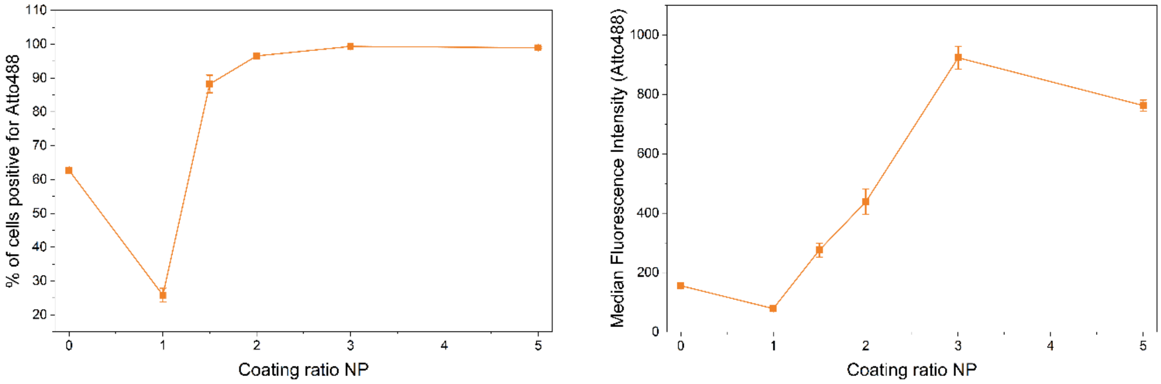

**Figure SI4. Uptake of PEG5K-K10 coated 24HB-F in ARPE-19 cells depending on the NP coating ratio with flow cytometry.** On the left, expressed as percentage of cells positive for Atto488 from the relevant single cell population. On the right, expressed as median fluorescence intensity of Atto488.

Similar to our observations with Y-79 cells, the increasing the coating ratio lead to enhanced cell association. Almost 100% Atto positive cells were achieved with an NP2 coating ratio. This could be either due to the enhanced phagocytic activity of ARPE-19 cells or because the 24HB-F has double as many attached Atto488 fluorophores compared to 24HB-RFLP that lead to a stronger fluorescence signal per cell, even though ARPE-19 cells were treated only with 1 nM of DONs (instead of 1.5 nM). The main difference to Y-79 cells was that uncoated 24HB-F showed an over 60% positive cell population that then dropped to under 30% at NP1. Considering the trend visible in the graph “Median fluorescence intensity vs. Coating ratio NP”, it would seem that the MFI strongly increased from NP1.5 onwards, suggesting that perhaps uncoated 24HB-F (NP0) were either just attached to the cells, or that low coating ratios like NP1 hinder uptake compared to plain 24HB-F, till they reach a certain charge threshold e.g. here around NP1.5, that promoted cell uptake. Previously with ARPE-19 cells,[3] we observed only limited cell uptake of 24HB-F, thus some sort of DON aggregation or attachment to the cell appears likely.

We also imaged ARPE-19 cells that were treated with uncoated and coated 24HB-F at NP ratios 1 and 2 with the confocal microscope (see Figure SI9). The outline of ARPE-19 cells was difficult to make out, but coated DONs showed clear distribution at the cell interface and possibly even breached the cytosol. Especially at NP2, we observed colocalization with Lysotracker DeepRed dye. For uncoated particles, the signal in the Atto488 channel showed more agglomerates, that partly could be in the cells but likely were attached to the outside of cells as well. Also, no colocalization with Lysotracker or even signal from the cytosol was visible and they seemed to be rather distributed throughout the sample. This was unexpected, as previously uncoated 24HB-F attached evenly around the cells without agglomeration,[3] possibly indicating some interaction between FOB, media components and the 24HB-F, that would require further exploration.

### SI8. Effect of PEG5K-K10 coating at different NP ratios on the uptake of coated 24HB DONs into Y-79 cells

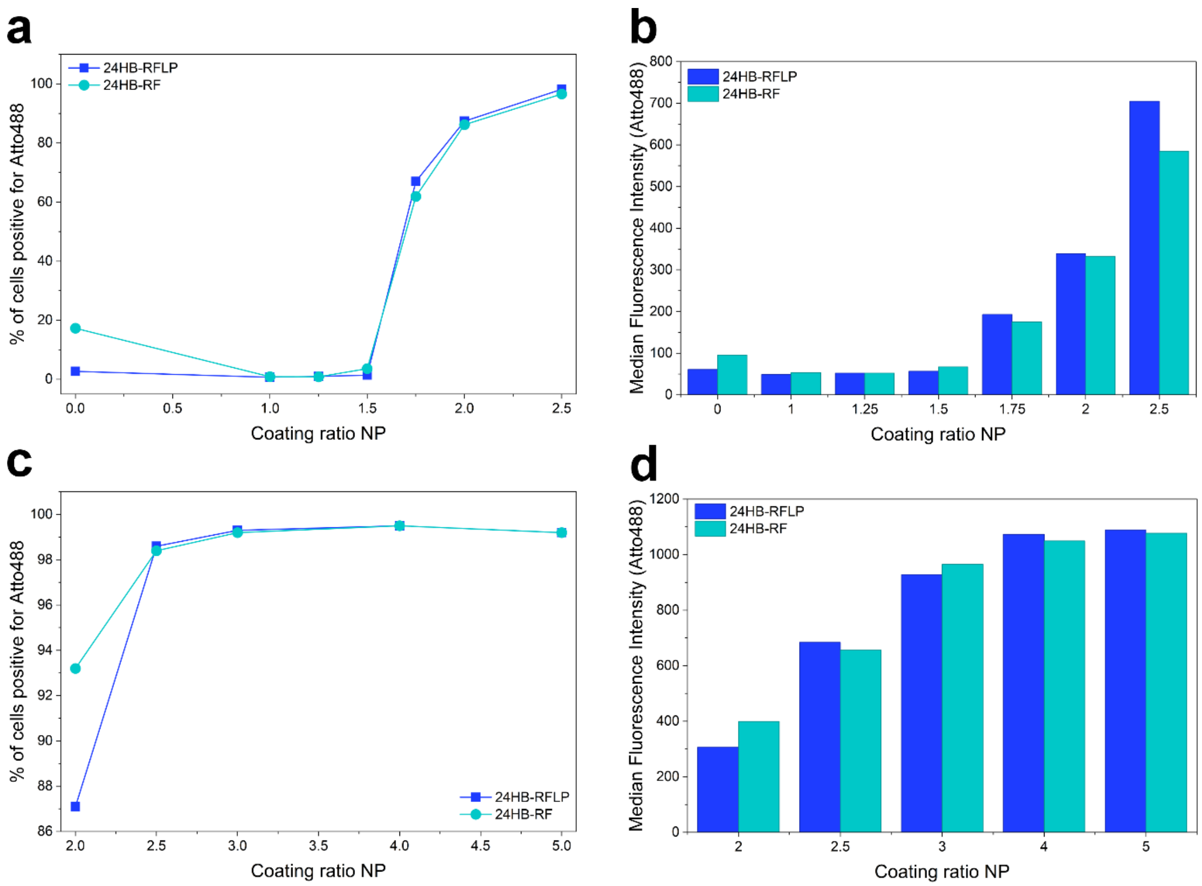

**Figure SI5. PEG5K-K10-coated 24HB-RFLP and 24HB-RF in Y-79 cells depending on the NP coating ratio with flow cytometry.** DONs in 1×FOB at 1.5 nM final concentration were incubated with Y-79 cells for 24 h at 20%(V/V) proportion in RPMI-1640 media with 1%PS. Percentage of cells positive for Atto488 after treatment with **a.** NP0-2.5 and **c.** NP2-5 coated DONs and their respective median fluorescence intensities (**b, d**). Each data point, corresponding to one single sample (no replicates), was plotted in dependence of the NP ratio to observe trends.

It became apparent that there seemed to be a specific threshold at coating ratio NP1.75 that started enabling cell uptake. At NP3, almost all cells gave an Atto488 signal. Increasing NP ratios raised the cell fluorescence intensity till it started leveling out around NP4 and 5. No prominent differences between coated 24HB-RF and 24HB-RFLP were observed, indicating these effects were mediated by the coating.

### SI9. Molecular dynamics simulation of DNA strands with PL3 peptides

All molecular dynamics (MD) simulations were performed using GROMACS (version 2025.2).[9] The CHARMM36m force field was employed to describe both peptides and DNA.[10] The TIP3P water model, compatible with the selected force field, was used to represent explicit solvent molecules.[11]

The simulated system consisted of a double-stranded B-DNA duplex (sequence: CGCGAATTCGCG) and five cysteine-PL3 peptide molecules (sequence: CAGRGRLVR). The initial DNA structure was obtained from 3DNA 2.0 web server,[12] and peptide structures were generated in extended conformations using the Avogadro modelling software prior to system assembly.[13] To enable unbiased observation of spontaneous binding events, the peptides were initially positioned randomly around the DNA molecule. The system was placed in a cubic simulation box with the dimensions of 7 × 7 × 7 nm and solvated with explicit water molecules. Counterions were added to neutralize the total system charge, and additional KCl was introduced to achieve a physiological salt concentration of 0.15 M.

Energy minimization was first carried out using the steepest descent algorithm[14] until the maximum force on any atom was below 1000 kJ $mol^{-1}$ $nm^{-1}$, ensuring removal of steric clashes and unfavorable contacts. This was followed by production simulations up to 600 ns in the NPT ensemble at 310 K and 1 bar using isotropic pressure coupling with the V-rescale thermostat[15] and Parrinello–Rahman barostat,[16] respectively. Long-range electrostatic interactions were treated using the Particle Mesh Ewald (PME) method[17] with a real-space cutoff of 1.2 nm. The same cutoff distance was applied for LJ interactions, with force-switching applied from 1.0 nm. All covalent bonds involving hydrogen atoms were constrained using the LINCS algorithm,[18] which enabled the use of a 2 fs integration time step. Periodic boundary conditions were applied in all three spatial dimensions. To ensure reproducibility and statistical robustness of the observed binding events, three independent replicates were carried out using different initial velocity distributions generated according to a Maxwell–Boltzmann distribution at 310 K.

Peptide–DNA interactions were characterized by calculation of number of contacts as a function of time utilizing gmx mindist program distributed with the GROMACS simulation package.[9] The cut-off for contacts was defined as 0.6 nm. During the simulations, peptides progressively approached and associated with the DNA molecule (Figure SI6). By the end of the 600 ns simulations, stable peptide–DNA complexes were observed in all independent replicates. In several trajectories, multiple peptides wrapped around the DNA helix and remained associated for prolonged periods, indicating the formation of stable complexes.[19]

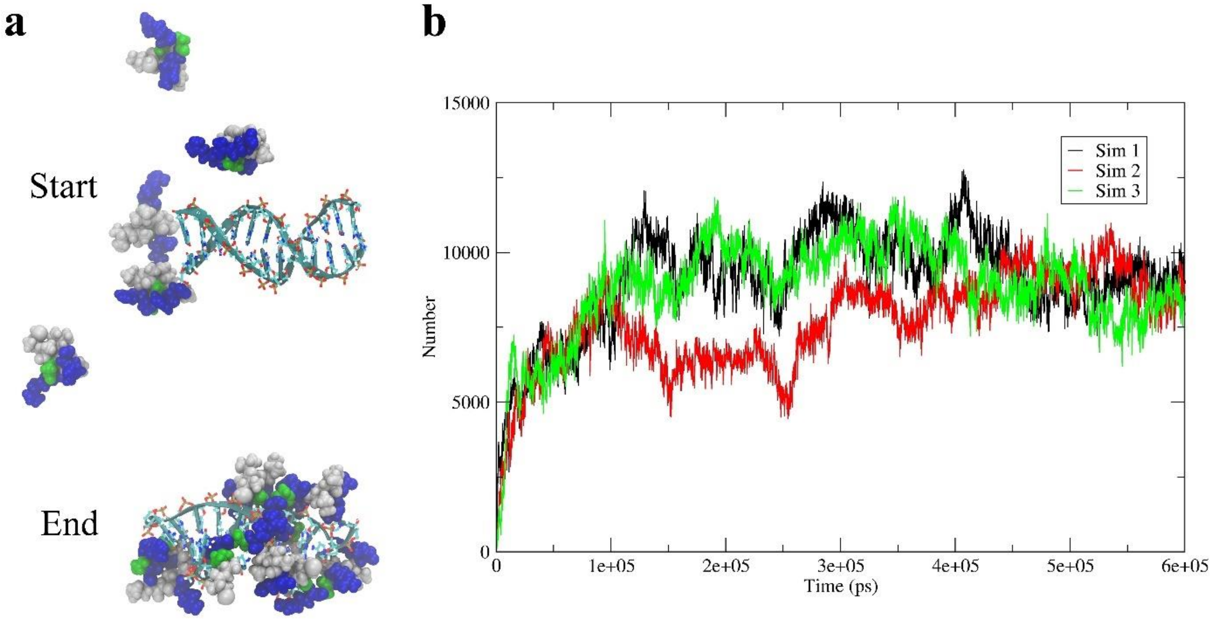

**Figure SI6. MD simulation of DNA and PL3 peptide interaction**. **a.** Representative snapshots from one simulation trajectory showing the initial and final configurations of the system. At the start of the simulation, peptides (green and white spheres represent polar and nonpolar residues whereas blue spheres highlight positively charged residues) were randomly distributed around the DNA duplex (cyan). During the simulation, peptides progressively approached and associated with the DNA. By the end of the 600 ns simulation, all peptides were bound and partially wrapped around the DNA helix, forming a stable peptide–DNA complex. **b.** Time evolution of the total number of peptide–DNA contacts as a function of simulation time for three independent replicates. In all simulations, the number of contacts increased rapidly during the early phase, followed by fluctuations around a plateau, indicating stable complex formation.

### SI10. AGE for specific incorporation of PL3-oligonucleotides into 24HB

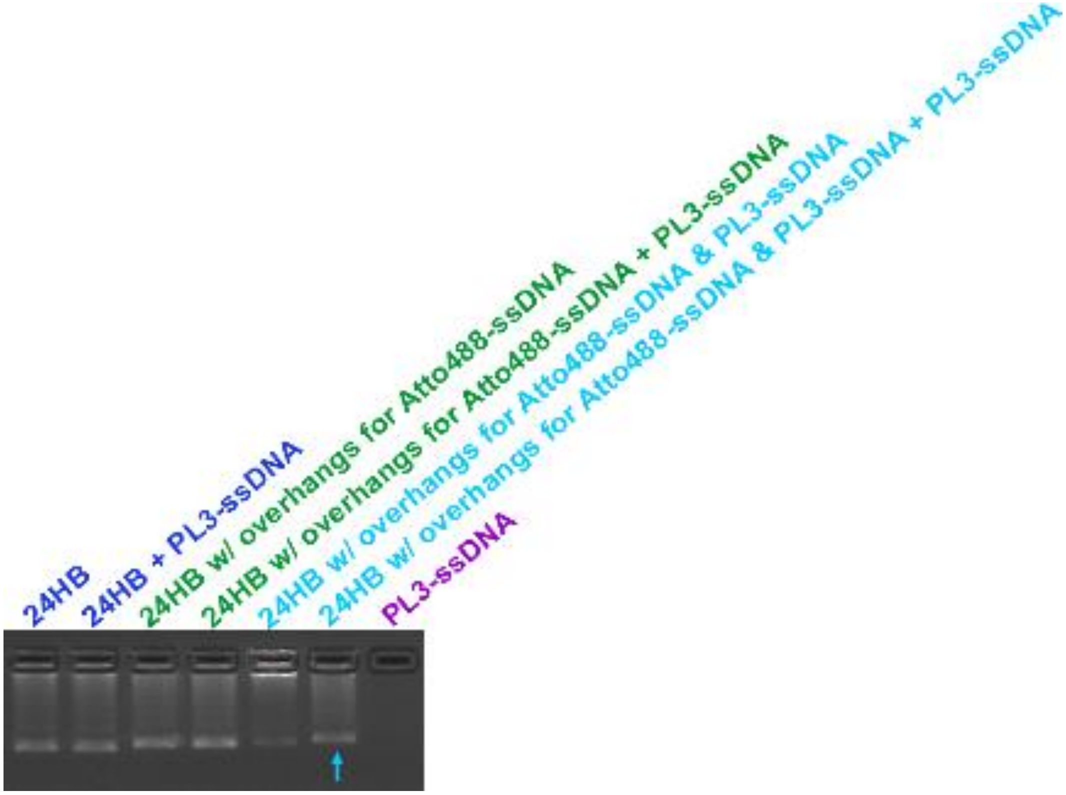

**Figure SI7. AGE check for unspecific incorporation of PL3-peptides.** To investigate unspecific, electrostatic attachment of PL3-ssDNA strands to 24HB DONs, PL3-oligonucleotides were incubated overnight with variations of 24HB DON that do not have any (24HB) or not the correct sequence-complimentary ssDNA strand extensions to hybridize PL3-oligonucleotides (24HB with overhangs for Atto488-oligonucleotides). The annealing reaction was prepared with a 10× excess of PL3-ssDNA per available attachment site and was incubated overnight at room temperature. Samples were diluted with 6×loading dye and run on a 2% agarose gel (90 V, 50 min on ice). The gel was imaged in the ethidium bromide channel.

For 24HB and 24HB with overhangs for Atto488-ssDNA, the coincubation with the PL3-ssDNA strands did not result in any shift of the band, while for 24HB with overhangs for Atto488-ssDNA and PL3-ssDNA a slight shift occurred (arrow). This indicated that stable peptide attachment to the 24HB structure was mediated by the specific PL3-ssDNA strands that hybridized to the base-complimentary overhang strands. While this cannot clarify how exactly the peptide is displayed on the DONs, it gave indication that electrostatic interaction alone would not lead to this observed band shift. Free PL3-ssDNA strands were run as additional control on the gel but did not give any signal in the ethidium bromide channel.

### SI11. Considerations for PEG conformation and coating thickness

As previously demonstrated and outlined by Rodríguez-Franco *et al.*,[8] we estimated the grafting density of our PEG5K-K10 coating at NP2 and PEG conformation as follows:[20–22]

**Estimation of surface area for 24HB:**

(simplified as cylindrical structure with approximate dimensions of 12 nm diameter by 115 nm length)

$$surface\ area = (2 * \pi * radius * height) + (2 * \pi * radius^2)$$

$$surface\ area = (2 * \pi * 6\,nm * 115\,nm) + (2 * \pi * (6\,nm)^2) \approx 4562\,nm^2$$

**Estimation of grafting density**:

(Values from Section SI2 were used)

One 24HB-RFLP particle has 16375 negative charges, meaning for a coating ratio of NP2, 32750 positive charges (provided by the polymer) are needed. Given that according to the certificate of analysis for our batch of PEG5K-K10, the degree of polymerization (number of monomeric units) of the poly-lysine block by NMR was 9, this results in around 3639 polymer molecules complexing with one 24HB-RFLP. That yields:

$$grafting\ density = \frac{number\ of\ polymer\ molecules}{surface\ area\ of\ 24HB} = \frac{3639}{4562\,nm^2} \approx 0.80\,polymer/nm^2$$

**Calculation Flory radius ($R_F$) and grafting distance (D)**[20–22]

Length of PEG monomer is 0.35 nm,[21] according to the manufacturer information PEG5K-K10 has 114 PEG units per polymer molecule, area that one PEG chain occupies is equal to the inverse of the grafting density (thus, for a grafting density of 0.80 polymer/nm$^2$ equals 1.25 nm$^2$/polymer)

$$R_F = length\ of\ monomer * \ number\ of\ PEG\ monomers\ per\ chain^{\frac{3}{5}} = 0.35\,nm * 114^{\frac{3}{5}} \approx 6.0\,nm$$

$$D = 2 * (\frac{area\ that\ one\ PEG\ chain\ occupies}{\pi})^{\frac{1}{2}} = 2 * (\frac{1.25\frac{nm^2}{PEG\ chain}}{\pi})^{\frac{1}{2}} \approx 1.26\,nm$$

**Conformational regime** was determined based on the **$R_F$/D ratio**:

$$\frac{R_F}{D} = \frac{6.0\,nm}{1.26\,nm} \approx 4.76$$

Since the $R_F/D$ ratio was larger than 1, the formation of PEG brush conformation (instead of a PEG mushroom) is assumed.[21–23] The maximum length of the brush would correspond to the length of monomer (0.35 nm) by the number of monomers (114), meaning in this case 39.9 nm. The actual thickness of the PEG layer thus presumably falls in between 6 nm and 39.9 nm. Some literature values on the thickness of PEG layers have been compiled and reported previously,[21,24] though because partially calculated or determined with various methods, showing variance and not exactly matching in conditions.

The oligonucleotide overhang strand used to attach PL3-oligonuclotide to our 24HB nanostructure is ~7.8 nm in length, so in combination with the higher coating ratio of NP2 that resulted in a high grafting density, there is a possibility that the targeting peptide might have been partially occluded or crowded by the PEG chain.

#### SI12. Live confocal images of Y-79 cells treated with NP2-24HB-RF

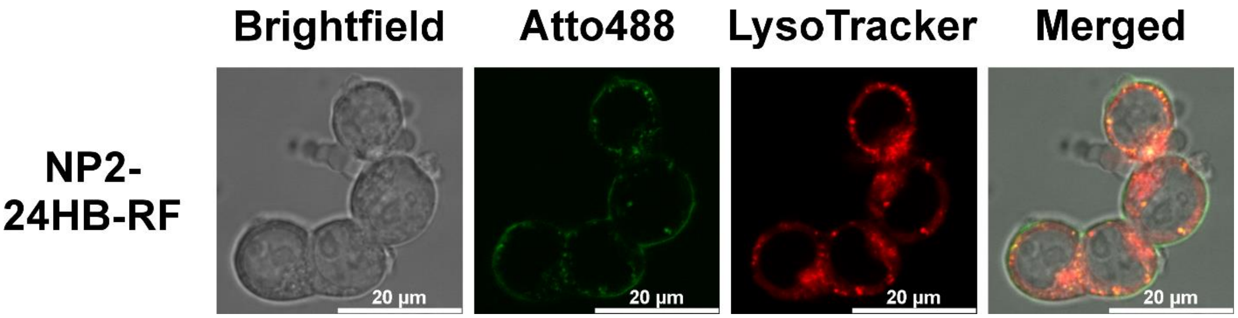

**Figure SI8. Uptake of NP2-24HB-RF in Y-79 cells.** Live confocal imaging of Y-79 cells treated with NP2-24HB-RF at 10 nM concentration in RPMI-1640 media with 1% PS after 24 h. 24HB-RF was functionalized with Atto488 fluorophores (green) and lysosomes in cells were stained with LysoTracker Deep Red (red). Colocalization of Atto488 and LysoTracker Deep Red results in yellow signal in the merged images.

### SI13. Live confocal images of coated 24HB DONs in ARPE-19 cells

#### Live confocal imaging for uptake study (ARPE-19)

10 000 ARPE-19 cells (100 µL/well) were seeded overnight to the CellView cell culture slide (TC, glass bottom, Greiner, 543078) and let attach at 37 °C with 5% $CO_2$. The following day, the FBS-containing CM was removed, the wells washed with 100 µL of 1×DPBS and 125 µL of DON treatments (in 1×FOB) in CM 1%PS (DMEM/F12, 1% PS) were added, resulting in a final DON concentration of 10 nM in a 20% (V/V) proportion with CM 1%PS. The cells were incubated at 37 °C with 5% $CO_2$ for 24 h. For the last hour of the incubation, LysoTracker Deep Red (Invitrogen) at a final concentration of 100 nM was co-incubated where needed. The imaging was performed similarly, as described in the main text for Y-79 cells.

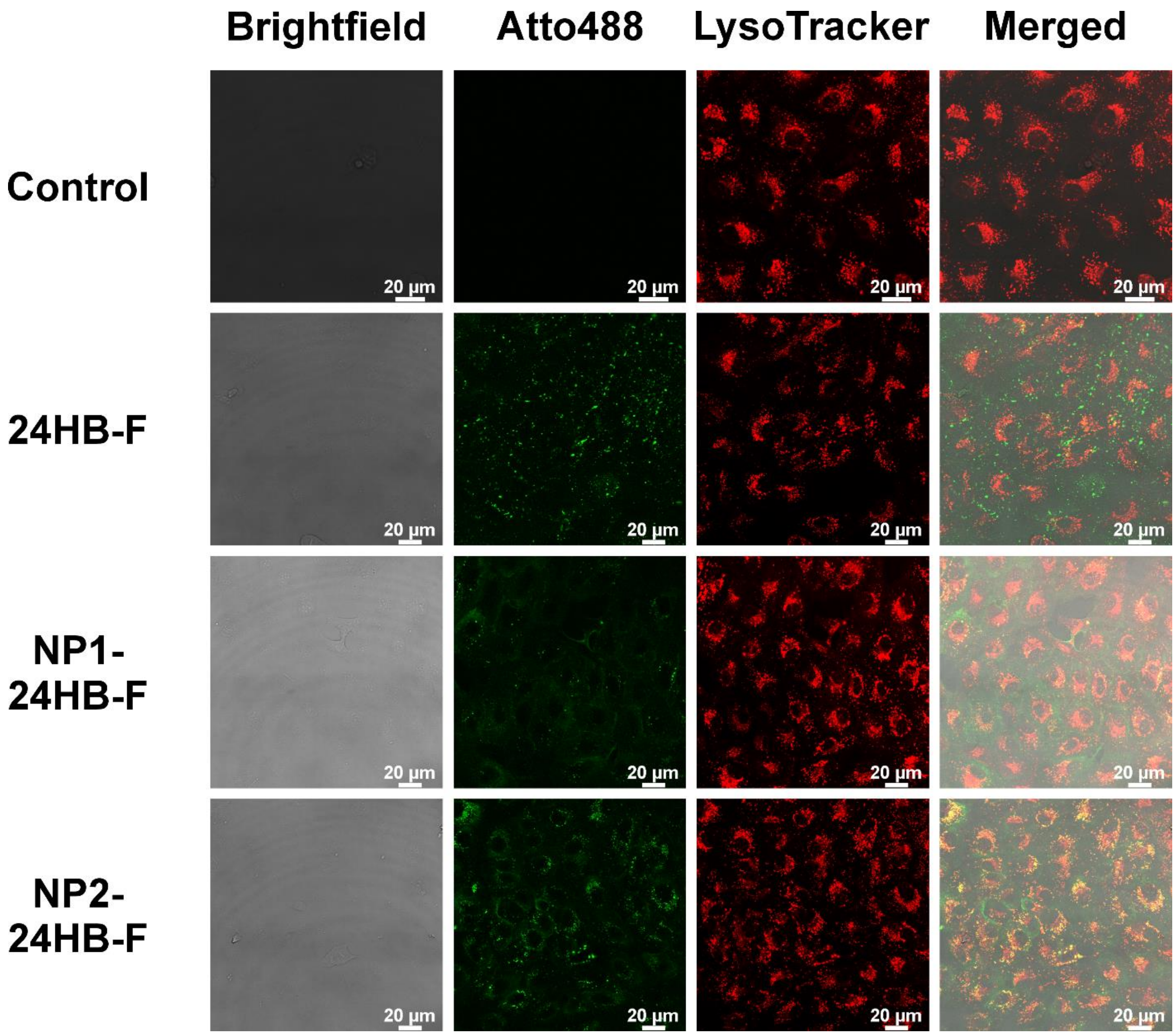

**Figure SI9. Live confocal imaging of ARPE-19 cells after treatment with uncoated and coated 24HB-F for 24 h.** DONs in 1×FOB were added at a final 10 nM concentration in a

20%(V/V) proportion in DMEM/F12 media with 1% PS. 24HB-F was functionalized with Atto488 fluorophores (green), lysosomes in cells were stained with LysoTracker Deep Red (red), buffer-treated cells served as control. Colocalization of Atto488 and LysoTracker Deep Red resulted in yellow color in the merged images.

Considering the more intense and frequent overlap of the Atto488 signal with Lysotracker, NP2-coated particles appeared to be more efficiently internalized compared to NP1-coated particles, in line with our observations from flow cytometry. The outline of ARPE-19 cells was hard to discern, but it seemed that coated DONs were not only present in the lysosomes but perhaps also in the cytosol or at least spread along some cell membranous structures. In contrast to Y-79 cells, they did not as strongly accumulate at the outer cell membrane. For uncoated 24HB-F, agglomerates were observed that in part were probably internalized but most likely were also attached on the outside of the cells, as they did not colocalize with Lysotracker and seemed randomly distributed throughout the sample. This was unexpected, as previously uncoated 24HB-F attached evenly around the cells without agglomeration.[3] This could indicate some interaction between FOB, media components and the 24HB-F, though it was not observed for any of the coated DONs, and would require further exploration.

### SI14. Cell viability assessment of PEG5K-K10 polymer and coated 24HB DONs in ARPE-19 cells (exploratory)

#### Cell viability assessment with alamarBlue Assay (ARPE-19)

In 96-well plates, 10 000 ARPE-19 cells were seeded in 100 µL/well of FBS-containing cell media and grown overnight at 37 °C with 5% $CO_2$. Free PEG5K-K10 (final concentrations: 0.5-100 µM) was tested in FBS-containing conditions, while PEG5K-K10-coated 24HB (final DON concentration: 2 nM or 10 nM) were assessed in FBS-free conditions. To this aim, the CM was removed for the relevant wells, the wells then washed once with 1×DPBS and replaced with 100 µL of CM 1%PS (DMEM/F12, 1% PS). 25 µL of treatments in 1×FOB were spiked into the wells at a final proportion of 20% (V/V) and incubated at 37 °C with 5% $CO_2$ for 24 h. Subsequently, treatments were removed, the wells washed with 1×DPBS, replaced with 150 µL of 1×alamarBlue HS Cell Viability Reagent (Invitrogen, A50100) in CM 1%PS and then incubated at 37 °C with 5% $CO_2$ for another 3.5 h. 100 µL of each well was transferred to a black-walled read-out plate (Thermo Scientific, 265301). The fluorescent measurements were performed with Varioskan LUX (Thermo Scientific, top optics, $\lambda_{exc}$= 560 nm, $\lambda_{em}$= 590 nm). This exploratory experiment was only performed once (n=1) with each treatment condition in technical triplicate. The cell viability was expressed as % viability relative to the buffer-treated control.

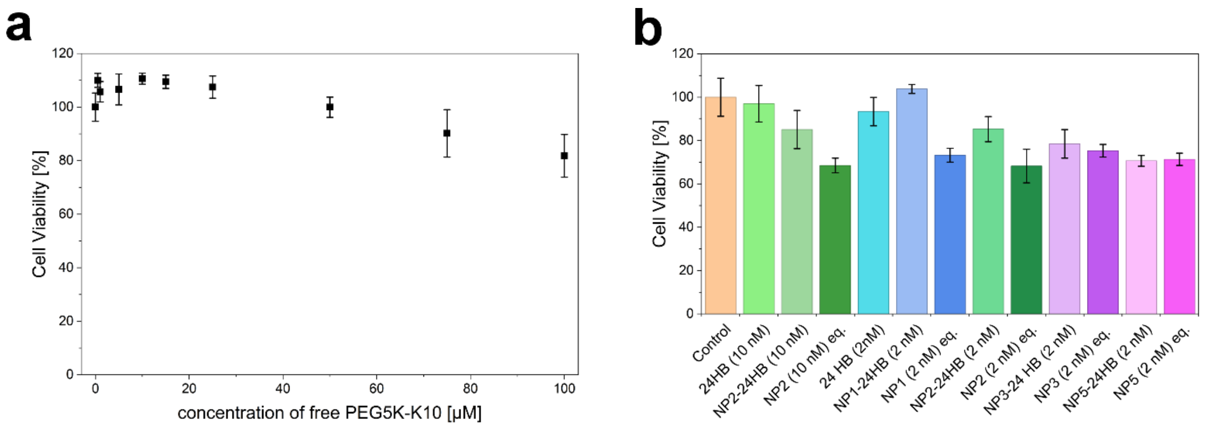

**Figure SI10. Cell viability of ARPE-19 cells after 24 h exposure, assessed *via* alamarBlue assay.** Cells were treated with **a.** free PEG5K-K10 polymer (0.5-100 µM) or **b.** uncoated (NP0) and coated 24HB (2 and 10 nM) at different coating ratios (NP1-5) and controls of corresponding amounts of free polymer (NPx equivalent (eq.)). ARPE-19 cells in **a** were exposed under FBS-containing conditions, while cells in **b** were treated under FBS-free conditions, in correspondence to the uptake experiments. Buffer-treated cells served as control (set to 100% viability). Mean ± s.d. was reported, acquired from one experiment (n=1) with three technical replicates.

In comparison to Y-79 cells, we observed little effect of free PEG5K-K10 polymer in this concentration range on the cell viability of ARPE-19 cells. There was a slight reduction (~20-30%) observed for coated DONs in accordance with the NP ratio. However, considering that these values were derived only from one single experiment, and that we have previously observed easily ±20% cell viability variations for ARPE-19 cells on different days and wells,[3] we believe that further replication is needed to verify such effects. Tentatively, we conclude that PEG5K-K10 polymers are well tolerated by ARPE-19 cells, given that free polymer did not show large toxic effects and that there was always little difference in viability between coated DONs and the equivalent free polymer. The only exception was NP1-24HB and its equivalent which would seem rather surprising since this corresponded to the lowest tested polymer amount. This would rather indicate a sensitivity to free polymer if not complexed with DONs which was however not observed for other samples. Furthermore, toxicity did not scale in a DON concentration-dependent manner, as NP2-24HB showed similar viability effects both at 2 nM and 10 nM concentration.

### SI15. Cell viability assessment of 24HB and NP2-24HB (10 nM) in Y-79 cells

**Table SI3. Cell viability of Y-79 cells after 24 h exposure, assessed *via* alamarBlue assay.** Cells were grown adherently and treated under FBS-free conditions with uncoated (NP0) and NP2-24HB in 1×FOB at 20%(V/V) proportion at 2 nM and 10 nM final concentration. Buffer-treated cells served as control (set to 100% viability). Mean ± s.d. was reported, acquired from three independent experiments (n=3) with each three technical replicates.

| Sample | Mean cell viability [%] | ± Standard deviation |
|---|---|---|
| 24HB (2 nM) | 115.7 | 10.5 |
| 24HB (10 nM) | 116.6 | 12.3 |
| NP2-24HB (2 nM) | 107.8 | 9.8 |
| Free polymer equivalent to NP2-24HB (2 nM) | 81.7 | 5.6 |
| NP2-24HB (10 nM) | 107.4 | 7.5 |
| Free polymer equivalent to NP2-24HB (10 nM) | 15.7 | 5.7 |

Uncoated and NP2-coated 24HB did not affect the cell viability of Y-79 cells at 2 nM and 10 nM concentration.

### SI16. Examples of trajectories during single particle tracking

Uncoated

NP2

NP5

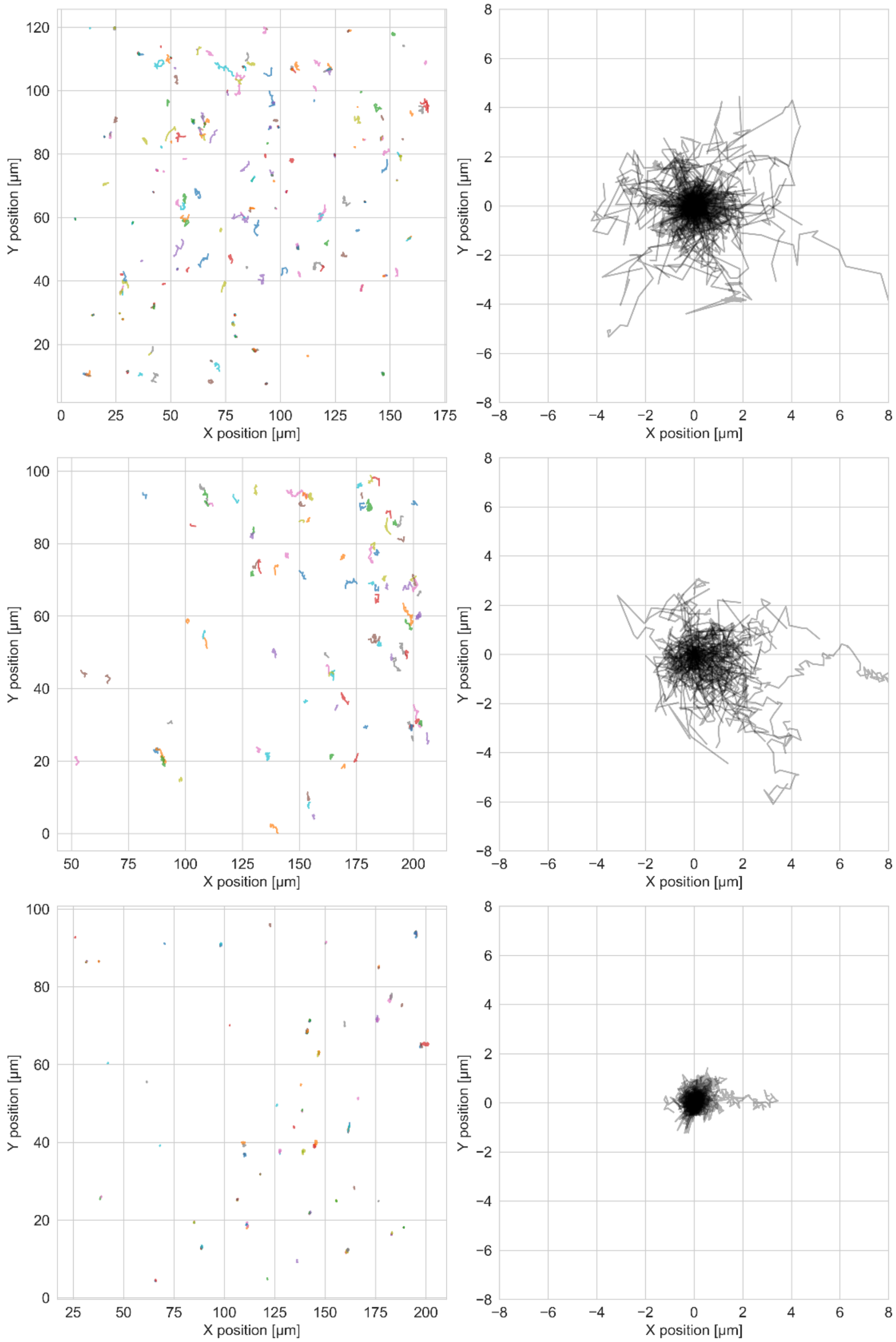

**Figure SI11**. **Exemplary trajectories during single particle tracking in *ex vivo* porcine eyes for uncoated and coated 24HB-F at NP2 and NP5 ratio (left) and same trajectories visualized with combined starting point (right).** Shorter and more condensed trajectories indicate less particle mobility.

### SI17. Stability of PEG5K-K10 coated 24HB DONs in porcine vitreous

#### Extraction and preparation of homogenized porcine vitreous

Porcine eyes were acquired and cleaned as described in the main methods section. Upon removal of the anterior part of the eye, the vitreous was collected and homogenized on ice with a glass tissue homogenizer. Subsequently, the vitreous was centrifuged at 4 °C (3200 g, 1 h) and decanted into a new falcon to remove melanin. The vitreous was filtered through a 0.45 µm and 0.22 µm sterile filter (Sartorius, 16537 and K16532 K) before storing aliquots at -80°C.

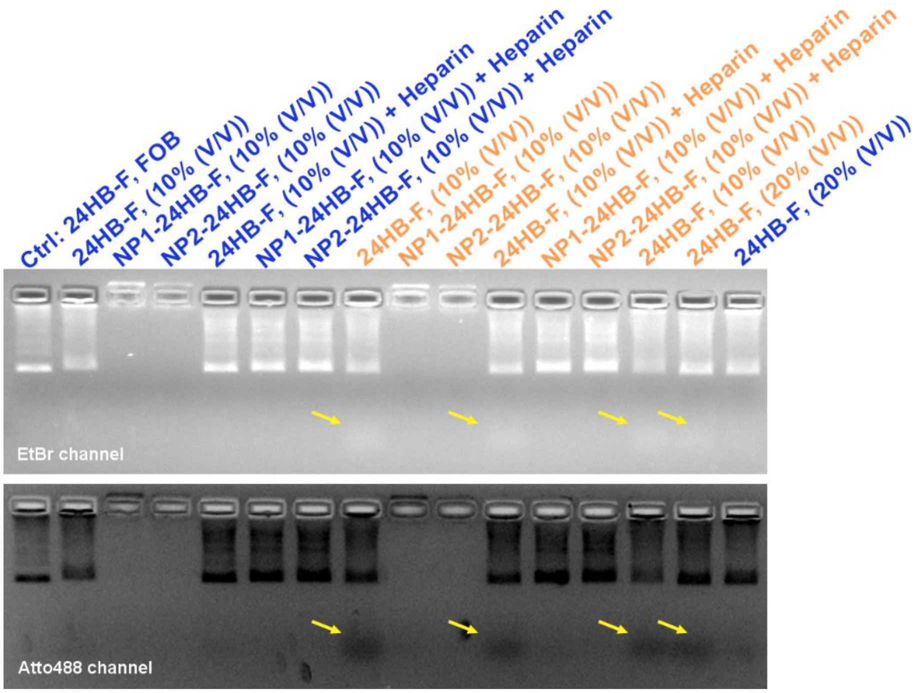

**Figure SI12. Stability of uncoated and PEG5K-K10-coated 24HB-F in homogenized porcine vitreous for 24 h at 4 °C (blue) and 37 °C (orange).** Incubation reactions were prepared at 3 nM concentration to a final volume of 20 µL, by mixing 24HB-F in 1×FOB (making up either 10% or 20% (V/V)) with homogenized porcine vitreous that was prior thawed on ice. After the incubation, the polymer coating was removed if desired with a 50× charge excess of a 5 µL heparin solution for 1 h at room temperature, while other samples were just diluted with water. For uncoated, heparin-treated 24HB-F controls, the heparin quantities were chosen equivalent to the quantities needed for a NP2 coating. Subsequently, the Mg concentrations were adjusted for all samples with 5 µL of $MgCl_2$/water to approximately match the Ctrl sample, without considering any preexisting $Mg^{2+}$ from the vitreous.[25] While this is not entirely accurate, bands for samples prepared at the same dilution proportion are comparable and no differences between the different bands were observed. Coated samples remained stably coated even in presence of the vitreous and at 37 °C. Uncoated 24HB-F showed some loss of the attached fluorophore at 37 °C, while no tag loss was observable for NP1 and NP2-coated 24HB-F, in line with previous observations that these coatings stabilize DONs under physiological conditions. Before starting the run on the 2% agarose gel, samples were diluted with 6×loading dye (90 V, 50 min on ice). Gels were imaged in both the ethidium bromide (EtBr) and Atto488 channel.

### SI18. Gating strategy for Y-79 cells in flow cytometry experiments

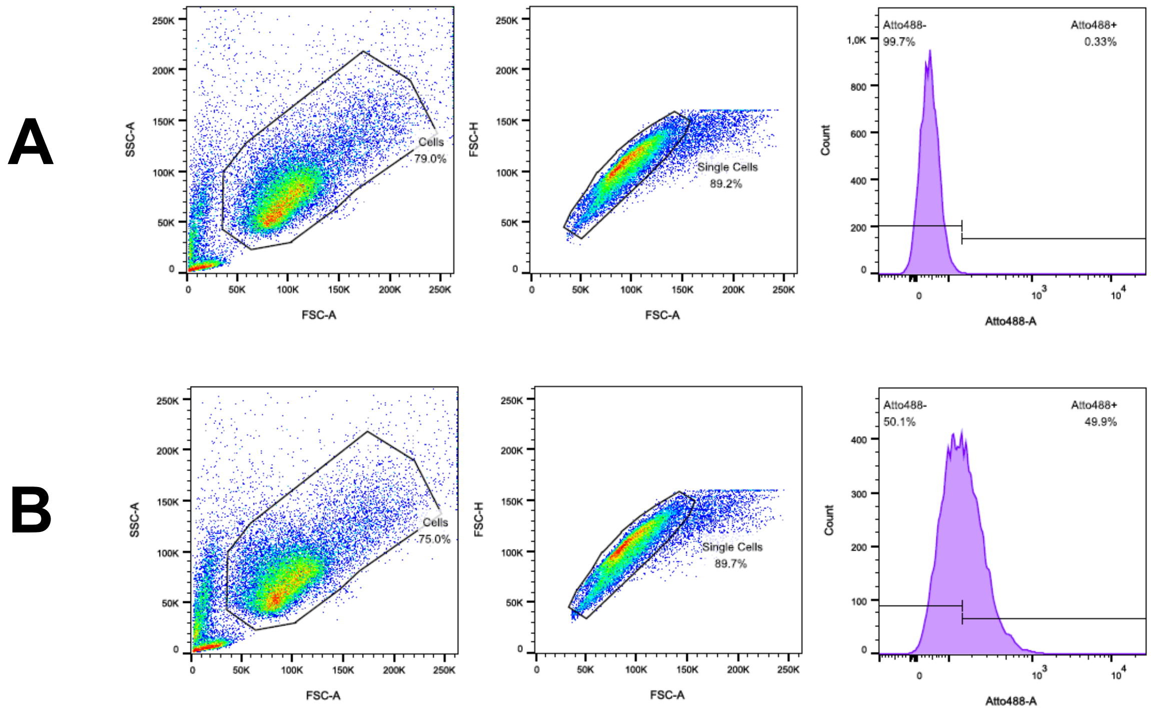

**Figure SI13. Representative gating strategy for Y-79 cells in flow cytometry experiments.** Row A shows buffer-treated control cells that were used to set the gate in the Atto488 channel for the max. 0.5% fluorescence overlap. Row B shows NP2-24HB-RFLP-treated cells (24 h). Plots from left to right: FSC-A vs. SSC-A (gating of live cells by size and granularity), FSC-A vs. FSC-H (doublet-discrimination), and the histogram for Atto488-A. Pseudocolour plots are shown, while contour plots were used additionally as guide for setting the gates.
